# Silk Fibroin Particles as Carriers in the Development of All-Natural Hemoglobin-Based Oxygen Carriers (HBOCs)

**DOI:** 10.1101/2023.03.01.530637

**Authors:** Marisa O Pacheco, Henry M Lutz, Jostin Armada, Nickolas Davies, Isabelle K Gerzenshtein, Alaura S Cakley, Bruce D Spiess, Whitney L Stoppel

## Abstract

Oxygen therapeutics have a range of applications in transfusion medicine and disease treatment. Synthetic molecules and all-natural or semi-synthetic hemoglobin-based oxygen carriers (HBOCs) have seen success as potential circulating oxygen carriers. However, many early HBOC products were removed from the market due to side effects from excess hemoglobin in the blood stream and hemoglobin entering the tissue. To overcome these issues, research has focused on increasing the molecular diameter of hemoglobin by polymerizing hemoglobin molecules or encapsulating hemoglobin in liposomal carriers, where immune responses and circulation times remain a challenge. This work looks to leverage the properties of silk fibroin, a cytocompatible and non-thrombogenic biopolymer, known to entrap protein-based cargo, to engineer a silk fibroin-hemoglobin-based oxygen carrier (sfHBOC). Herein, an all-aqueous solvent evaporation technique was used to form silk fibroin particles with and without hemoglobin to tailor the formulation for specific particle sizes. The encapsulation efficiency and ferrous state of hemoglobin were analyzed, resulting in 60% encapsulation efficiency and a maximum of 20% ferric hemoglobin, yielding 100 µg/mL active hemoglobin in certain sfHBOC formulations. The system did not elicit a strong inflammation response *in vitro*, demonstrating the potential for this particle system to serve as an injectable HBOC.

**Table of Contents Figure:** 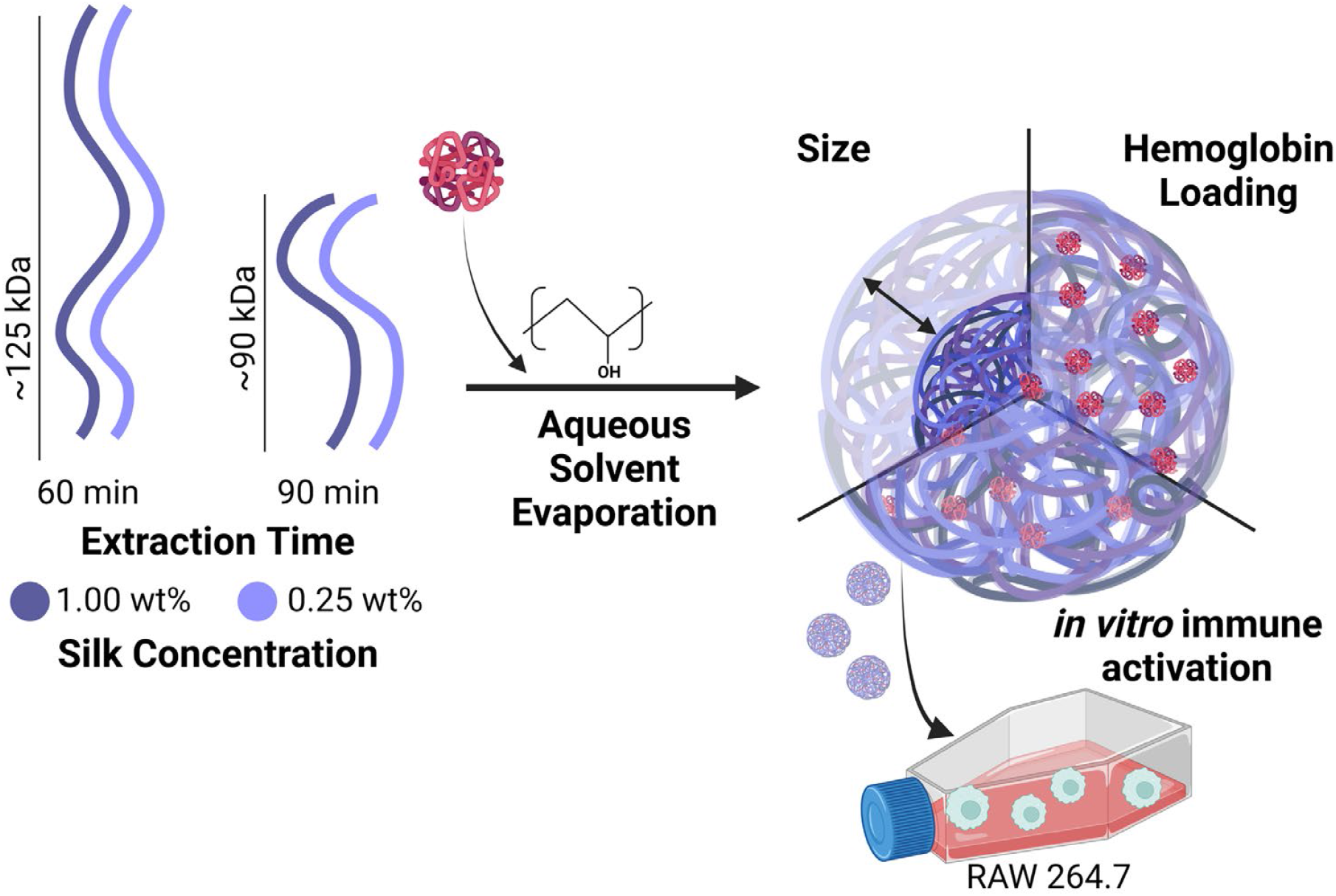

**Table of Contents Figure:** In this manuscript, we generate silk fibroin particles using an all-aqueous processing technique starting from silk fibroin polymer systems of differing molecular weights. We analyze the extent to which silk concentration and extraction time affect particle size. Further, we analyze the encapsulation of hemoglobin in the particle system and assess immune activation in macrophage-like cultures.

## 1. Introduction

Many injuries and diseases lead to local tissue hypoxia and/or systemic hypoxemia, with both acute and chronic diagnoses. Causes or co-morbidities include, but are not limited to, traumatic injury, sickle cell disease, diabetes, COVID-19, kidney disease, stroke, and other hematologic conditions. One treatment strategy is respiratory supplementation of oxygen to better saturate native red blood cells (RBCs). Another strategy is transfusing donated blood or blood components to improve systemic oxygen carrying capacity. Maintaining a commercial or clinical blood supply for treatment of diseases or conditions resulting in low oxygen tension remains a critical challenge. Blood transfusion and blood product supply chain issues result from factors such as the limited shelf-life of stored RBC units,^2, 3^ spikes in demand due to seasonal^4, 5^ or emergency surges,^6, 7^ and breakdown of the supply chain,^8^ motivating the need to develop alternatives to RBC transfusions that can overcome these dynamic challenges. The developed replacement ideally should replicate the oxygen carrying function of RBCs effectively and safely with focus on pathogen-free and universally-compatible solutions that can overcome transportation and storage limitations (see ^9^ for a recent perspective on this topic).

Over the past several decades, many materials, including synthetic perfluorocarbons^10^ and hemoglobin-based oxygen carriers (HBOCs) based on human,^11^ bovine,^12-14^ and recombinant hemoglobin,^15^ saw success in *in vitro* studies, early-stage *in vivo* models, and made it to phase I clinical trials. However, early HBOCs were found to cause severe side effects as a function of two main factors. First, it was determined that particle extravasation into the tissue space through the vessel wall (fenestration diameter ~50-100 nm^16^) caused scavenging of nitric oxide (NO), leading to systemic hypertension and oxidative tissue injury.^17^ Second, excess cell-free hemoglobin present in circulation overwhelmed the native clearance processes executed by macrophages in the liver and spleen, leading to renal toxicity.^18^ To address these challenges, strategies to increase the molecular diameter of the HBOC and limit free hemoglobin in circulation have become requirements in the design of modern HBOCs. These strategies include chemical crosslinking of hemoglobin,^19, 20^ polymerization of hemoglobin and crosslinked hemoglobins,^21-23^ and encapsulation of hemoglobin in liposome and nano- and microparticle systems.^24, 25^ Liposome encapsulated hemoglobins have demonstrated many of the desired design criteria for an alternative oxygen carrier, including absence of blood group antigens and blood-borne pathogens and a prolonged storage capacity.^26-28^

While liposome encapsulated systems show great potential for the eventual translation of new oxygen carrier technologies, several challenges still exist. One challenge is complex and expensive processes for preparation of liposomal systems^29^ as well as the use of harsh solvents and complex techniques often required for their synthesis.^30^ In this study we evaluate the encapsulation of hemoglobin in a silk fibroin particle system to limit the potential cost of synthesis^31^ while maintaining encapsulated hemoglobin structure and oxygen biding and release capabilities. Silk fibroin is a biopolymer isolated from the cocoons of *Bombyx mori* silkworms and in formats such as films, has demonstrated effective encapsulation and stabilization of large, bioactive proteins and other blood components^32, 33^ as well as biocompatible and antithrombogenic behavior.^34, 35^ Studies suggest that the semi-crystalline nature of the silk fibroin protein can be leveraged to stabilize the structure of proteins^36^ that are entrapped within the crystal structure of the β-sheet domains present in the heavy chain of silk fibroin. Utilizing this hydrophobic collapse of the silk fibroin protein allows for addition of these bioactive components in the soluble state, but stabilization in both hydrated and dry (lyophilized) states are possible (see reference for a deeper review^33^).

Bioactive molecule encapsulation and stability in silk fibroin nano- and microparticles has previously been investigated in the context of cancer nanomedicines and potential intraperitoneal delivery.^37-41^ Due to the nature of anti-cancer drugs being mainly small molecules and the complexity in studying natural biopolymer and small molecule interaction in a crystallized state, there is limited knowledge of how silk particle systems encapsulate and stabilize larger protein-based cargo. We also seek to understand if stabilization in the nano- or microparticle systems differs from the β-sheet crystalline driven process observed in macro-sized silk materials such as films and scaffolds.^33, 42^ Additionally, the most common method of synthesis for silk-based anticancer nanomedicines is organic solvent desolvation,^43^ which would potentially cause similar damage to hemoglobin cargo as do harsh liposome synthesis methods. Thus, we look to take advantage of the ability of silk fibroin biopolymers to stabilize protein cargo while leveraging an all-aqueous particle fabrication technique inspired by the synthesis of silk nano-porous films.^44, 45^

In this work, we encapsulate human hemoglobin in silk fibroin nano- and microparticles, focusing on the characterization of the parameter space available for making these particles with a focus on reproducibility and hemoglobin structure and loading efficiency. We aim to form nanoparticle formulations with a size distribution between 200-700 nm, with all nanoparticles being larger than 100 nm to avoid the potential for extravasation through the vessel wall. An ideal formulation would minimize the variability in particle size and show high reproducibility from batch to batch. To evaluate the potential for silk fibroin nanoparticles to stabilize and carry hemoglobin, we characterize the size, crystalline character, and internal morphology of four separate formulations chosen to modulate particle size. We then evaluate the passive encapsulation of hemoglobin at two concentrations in the same particle formulations. Lastly, we assess the cell-material interaction of our sfHBOC system with macrophage-like cells in vitro to better understand the potential safety and clearance of the sfHBOCs.

## 2. Results and Discussion

Encapsulation of hemoglobin in an all-natural protein carrier that ensures maintenance of hemoglobin structure and function within the biopolymer network has the potential to address current challenges in the field of hemoglobin-based oxygen carriers. In this work, we utilize silk fibroin proteins isolated from *Bombyx mori* silkworm cocoons as the stabilizing biopolymer and carrier system. We approach this development of a silk fibroin hemoglobin-based oxygen carrier wholistically, first characterizing the silk fibroin polymer solution itself and ending with the encapsulation of hemoglobin in the created nanoparticle system and its interaction with macrophage like-cells *in vitro*. Initial experiments evaluating formulation parameters for size and morphology were completed with a hemoglobin solution of primarily methemoglobin (non-O_2_ binding Hb, denoted as mHb), but subsequent experiments assessing binding of O_2_ and cell response utilized HbA_0_, a non-glycated, ferrous form of hemoglobin (Millipore Sigma, H0267). sfHBOCs are formed using an all-aqueous method where the silk biopolymer is phase separated and crystallized using a bulk solution of 5% polyvinyl alcohol. We start by characterizing the neat silk particle parameter space before introducing hemoglobin and forming sfHBOCs (**Table 1**).

### 2.1. Organic solvent-free methods yield silk particles in size range of current HBOCs

#### 2.1.1. Extraction time yields control of precursor silk polymer solution molecular weight ranges

In order to reduce variability in sfHBOC particle size, robust control over and reduction of variation in biopolymer molecular weight is needed. In the silk fibroin biomaterials field, it is well understood that the extraction time of the fibroin polymer impacts the final material properties regardless of material format, with trends in extraction time (15, 30, and 60 minutes) being inversely proportional to biopolymer molecular weight.^46^ Thus, we hypothesized that longer extraction times would not only lead to shorter average fibroin chain lengths, but also a less variable polymer solution. Using sodium dodecyl sulfate-polyacrylamide gel electrophoresis (SDS-PAGE, **Figure S1**), we measured silk fibroin molecular weights and confirmed trends in published literature^46, 47^ as shown in **Figure 1A**. We determined that 90-minute extraction times yielded the most uniform polymer solutions with the lowest molecular weights, compared to 15, 30, and 60 minutes of extraction (**Figure 1B, S1**). Moving forward, we will use the 60-minute and 90-minute extracted silk fibroin solutions for particle formulations. The 60-minute extracted solutions had a median molecular weight of 125 + 3 kilodaltons (kDa) while the 90-minute extracted solutions had a median molecular weight of 90 + 1 kDa. We determined that the silk fibroin polymer chain lengths are heterogenous as shown by determination of effective interquartile ranges (IQRs) for these silk solutions (**Figure 1A, S1**). We calculated an IQR of ~95-105 kDa for 60 minutes of extraction verses an IQR of ~60-65 kDa for 90 minutes of extraction. While this heterogeneity in the biopolymer solutions exists, we wanted to confirm that the heterogeneity was consistent from batch to batch in terms of both the median molecular weight and level of heterogeneity. To test our hypothesis, we analyzed four separate 60-minute extractions and found that the polymer chain length and distribution were consistent from batch to batch as analyzed in **Figure 1CD**. Taken together, the results suggest that making silk particles with 90-minute extracted silk may lead to smaller and more uniform particle distributions.

**Figure 1:**
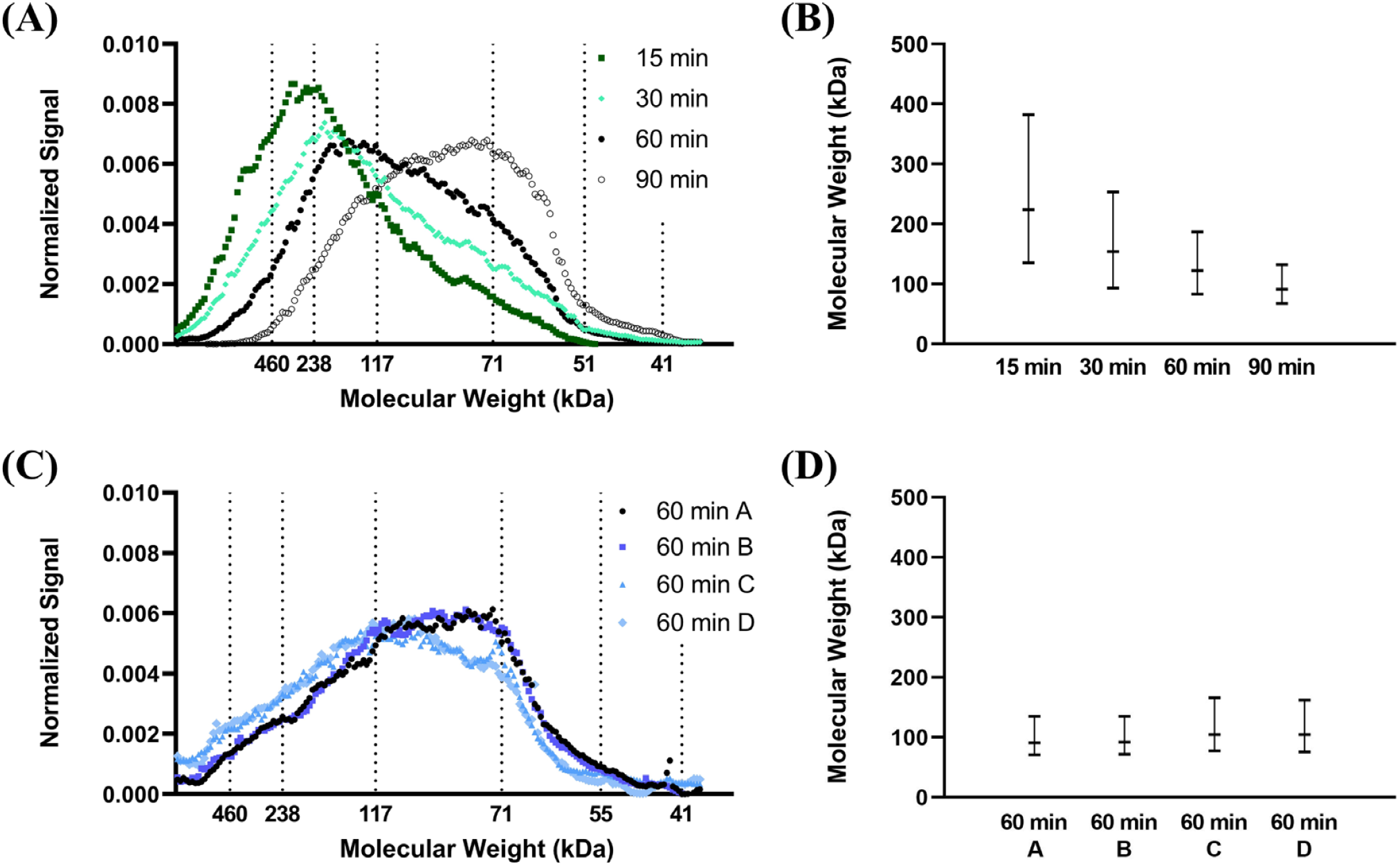
Control of silk polymer molecular weight. **(A)** Molecular weight distributions of silk polymer as a function of extraction time determined by analysis of SDS-PAGE gels. 2 µg of silk polymer was loaded in each lane of a 7% Tris-Acetate gel under reducing conditions. Densiometric analysis of colloidal blue stained gels yield the molecular weight distribution shown. **(B)** Effective interquartile range (IQR) of silk molecular weight as a function of extraction time. Horizontal data points represent the molecular weights at which the area under the distribution curve is equal to 25, 50, 75% of the total area. **(C)** Molecular weight distributions of 4 batches of silk polymer solution that was extracted for 60 min determined by SDS-PAGE. Densiometric analysis of colloidal blue stained gels yield the molecular weight distribution shown. **(D)** Effective interquartile range (IQR) of silk molecular weight as a function of batch. Horizontal data points represent the molecular weights at which the area under the distribution curve is equal to 25, 50, 75% of the total area.

#### 2.1.2. Decreased silk molecular weight and concentration yields smaller particles

For applications in delivery of hemoglobin via systemic injection of sfHBOCs into the blood, particles in the range of 200 nm - 1 μm are desired.^9^ In addition, for these particle systems to be applicable in a clinical setting, control over both particle size and distribution is required. Given that the 90-minute extracted silk fibroin solution has the lowest distribution of molecular weights, we hypothesized that utilizing 90-minute extracted silk fibroin solution in an all-aqueous silk particle fabrication method^44, 45^ would yield smaller and more monodisperse silk particles in solution. In addition, we recognize that the level of biopolymer entanglement during the phase separation process will dictate the theoretical number of polymers present in each particle and that previous work demonstrates the impact of polymer concentration on particle size.^48, 49^ Thus, we hypothesized that lower silk concentrations and lower silk molecular weights would lead to fewer secondary structure interpolymer interactions, resulting in decreased average silk particle size.

To determine the extent to which molecular weight and concentration impact particle size, we quantified particle hydrodynamic diameters using dynamic light scattering (DLS, **Figure 2AB**) and confirmed results with scanning electron microscopy (SEM, **Figure 2CDEF**). We compared particle formulations using 60- and 90-minute extracted silk fibroin at concentrations of 1% w/v and 0.25% w/v (**Figure 2A)**. Results show that extraction time, or the median molecular weight, is most impactful in dictating particle size. Particles formed at the two extremes, given by 1% w/v 60-minute extracted silk (60 min-1%-0 mHb, D_H_ = 490 + 70 nm) and 0.25% w/v 90-minute extracted silk (90 min-0.25%-0 mHb, D_H_ = 301 + 33 nm) are statistically different (p < 0.01) (see **Table 1** for all formulation descriptions). The 90 min-0.25%-0 mHb formulation maintains this statistical difference from the other evaluated formulations (p < 0.01, **Figure 2A**). It is important to note that the reported DH is the intensity weighted average hydrodynamic diameter determined by DLS, which can be inaccurate in the case of large, polydisperse samples. We performed additional analyses looking at the full intensity distribution functions and autocorrelation functions (**Figure S2 B-E**) Additionally, we calculated the difference between the intensity weighted average D_H_ and the hydrodynamic diameter that gives the maximum intensity (**Figure S2A**). In normal distributions with low to moderate polydispersity we expect this difference to be small, and we determined this was true for 90-minute formulations (**Figure S2A**) This led us to focus our analysis to the 90-minute formulations. In an assessment of dispersity, we calculated the full peak width at 68% (**Figure 2B**). We determined that lower concentrations led to tighter distributions, though the difference was not statistically significant. Qualitatively, SEM micrographs show that particle size decreases with decreasing biopolymer molecular weight, with quite observable differences at the extremes (**Figure 2C** compared to **Figure 2F)**. Additionally, SEM micrographs confirm the polydispersity we expected in the 60-minute conditions **(Figure 2CD**). Our results agree with those found in the literature, that modulation of silk particle size can be achieved using polymer concentration and molecular weight in the context of different fabrication strategies, such as microfluidic devices and organic solvent desolvation.^35, 37, 50, 51^

**Table 1:**
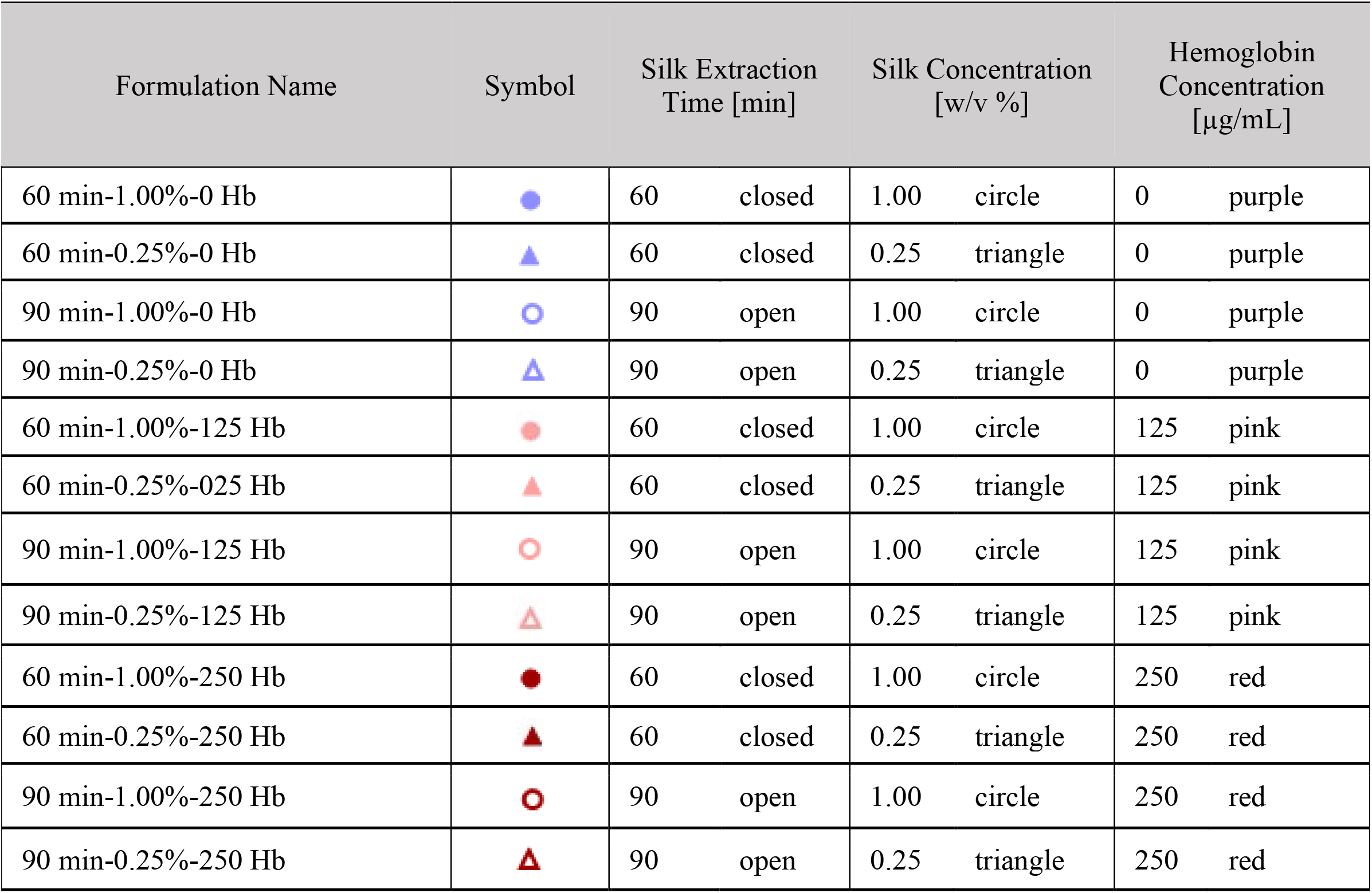
Formulation name, associated symbol, and details for particle formations used throughout studies presented in this manuscript.

| Formulation Name | Symbol | Silk Extraction Time [min] | | Silk Concentration [w/v %] | | Hemoglobin Concentration [$\mu\text{g/mL}$] | |
| --- | --- | --- | --- | --- | --- | --- | --- |
| 60 min-1.00%-0 Hb |  | 60 | closed | 1.00 | circle | 0 | purple |
| 60 min-0.25%-0 Hb |  | 60 | closed | 0.25 | triangle | 0 | purple |
| 90 min-1.00%-0 Hb |  | 90 | open | 1.00 | circle | 0 | purple |
| 90 min-0.25%-0 Hb |  | 90 | open | 0.25 | triangle | 0 | purple |
| 60 min-1.00%-125 Hb |  | 60 | closed | 1.00 | circle | 125 | pink |
| 60 min-0.25%-025 Hb |  | 60 | closed | 0.25 | triangle | 125 | pink |
| 90 min-1.00%-125 Hb |  | 90 | open | 1.00 | circle | 125 | pink |
| 90 min-0.25%-125 Hb |  | 90 | open | 0.25 | triangle | 125 | pink |
| 60 min-1.00%-250 Hb |  | 60 | closed | 1.00 | circle | 250 | red |
| 60 min-0.25%-250 Hb |  | 60 | closed | 0.25 | triangle | 250 | red |
| 90 min-1.00%-250 Hb |  | 90 | open | 1.00 | circle | 250 | red |
| 90 min-0.25%-250 Hb |  | 90 | open | 0.25 | triangle | 250 | red |

**Figure 2:**
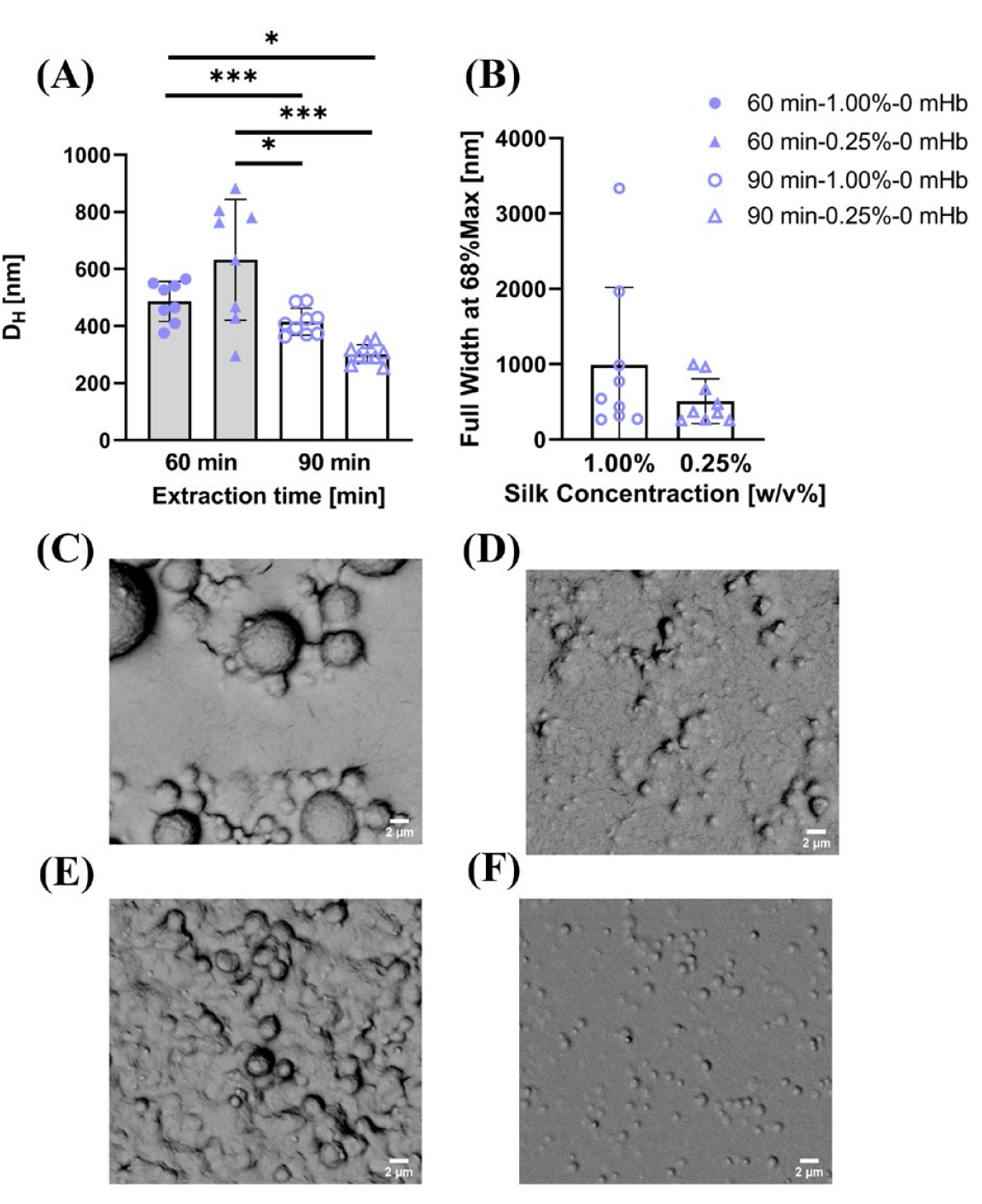
Silk molecular weight modulates particle size. (**A**) Comparison of hydrodynamic diameter (DH) for all silk only formulations as measured by DLS. (**B**) Comparison of peak width at 68% of the maximum, an assessment of dispersity. (**C**) Representative SEM micrograph for 60 min-1%-0 mHb (**D**) Representative SEM micrograph for 60 min-0.25%-0 mHb (**E**) Representative SEM micrograph for 90 min-1%-0 mHb (**F**) Representative SEM micrograph for 90 min-0.25%-0 mHb. It should be noted that the average silk particle hydrodynamic diameter is measured in a hydrated state, while SEM micrographs are captured after particles have been dried, and thus diameters estimated from SEM micrographs will not necessarily align with those determined by DLS. Data represented as AVG ± SD (n≥8). A two-way ANOVA with Tukey post hoc testing was performed. Statistical significance is shown as *p≤0.05, **p≤0.01,***p≤0.001, ****p≤0.0001.

To further assess structural changes in particles formed with different formulations, we analyzed crystalline content and internal particle structure using Fourier transform infrared (FTIR) spectroscopy and nanoscale computed tomography (Nano-CT), respectively. FTIR spectra can be compared to assess relative changes in crystalline content across formulations. This is done by deconvoluting the peak in the Amide I region^1^, shown in **Figure 3A**. The deconvolution allows for the determination of relative amounts of secondary protein structures present in the silk particles (**Figure S3**) and β-sheet or crystalline content is reported in **Figure 3B**. Results show silk concentrations impact total crystalline content, with a two-way ANOVA showing the variable of silk concentration is significant (p=0.0071). However, pair-wise comparisons showed no significance with average β-sheet content being around 30-40% in the formulations of interest (**Figure 3B**). High levels of crystalline content in our particle system is important for two reasons: (1) stable particles are formed by a hydrophobic collapse process, indicating that β-sheet crystalline structures should be present,^51-53^ and (2) bioactive molecule stabilization is expected to be achieved by kinetically trapping the molecules in crystalline structures.^33^ Thus, the consistency in high levels of crystalline content found in our particles is desirable for future applications, as the stabilization of the bioactive molecule, in this case hemoglobin, will be theoretically achieved evenly across formulations using 90-minute extracted silk.

**Figure 3:**
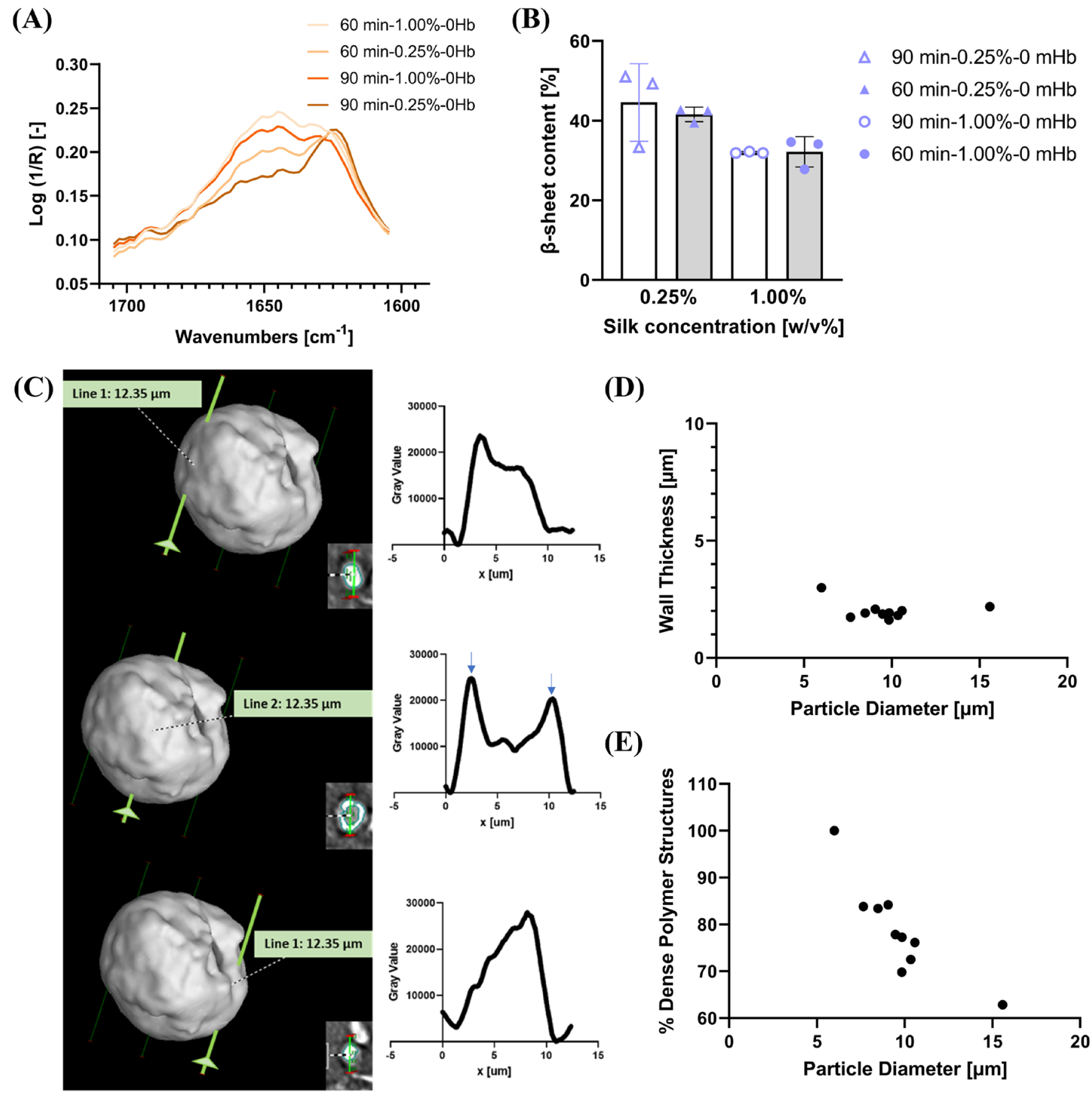
Highly crystalline domains are present across neat silk particle formulations. (**A**) FTIR spectra of the Amide I region for all neat silk formulations. Samples were dried and FTIR spectra were collected using a micro attenuated total reflections technique. (**B**) Following deconvolution of the amide I peak following published methods,^1^ relative level of β-sheet crystaillinity was determined. Data is represented as AVG ± SD (n=3). A two-way ANOVA with Tukey post hoc testing was performed. Statistical significance is shown as, *p≤0.05, **p≤0.01, ***p≤0.001, ****p≤0.0001 (**C**) Nano-CT was performed to observe if differences in crystallinity or polymer density could be observed within a single particle. Shown is a 3D reconstruction of a silk microparticle. Line profiles of the gray value from 3 2D Nano-CT slices (inset) for each green line are shown to the right. Blue arrows indicate dense wall structures. (**D**) Quantification of dense wall thickness via Nano-CT, shown is the thickness of the wall as a function of particle diameter for 10 particles imaged with Nano-CT. (**E**) The calculated percent of dense volume in a particle as a function of diameter assuming the particle is a sphere.

#### 2.1.3. Discussion of the role of crystallinity in protein encapsulation

As such, β-sheet or crystalline content of the particles is critical for understanding the potential to include and stabilize hemoglobin within these particles to create sfHBOCs. As previously hypothesized^33^, proteins and biomolecules are stabilized within the crystalline domains of the silk protein, trapping the bioactive molecule, in this case hemoglobin, in a stable confirmation even when the particles experience various hydration levels or temperatures. This feature of silk fibroin biomaterials is what makes this all-natural carrier system advantageous as a HBOC. It is important to recognize that FTIR spectra are representative of a combined particle population and cannot give information at the resolution of a single particle. Given that the bulk analysis of FTIR spectra did not identify significant differences in crystalline content, we further explored the internal structure of the silk particles to better understand the regions of potential hemoglobin incorporation. Due to the resolution of the Nano-CT, only particles on the micron scale can be adequately analyzed. Analysis of 10+ particles, ~5-15 μm in diameter, showed that the particles contained a denser wall structure with an inner spongy core (**Figure 3C**). The particles maintained a consistent wall thickness of 2 µm, suggesting that submicron particles are predominately dense polymer materials without a spongy core.

Together, results show that these formulations yield solid semicrystalline particles (**Figure 3DE**). The Nano-CT and FTIR results suggest that hemoglobin has the potential to be incorporated into the crystalline regions present throughout the particle, but that more hemoglobin may be captured in the denser outer wall due to changes in relative polymer density, in particles larger than 2 µm (outside the target size range of sfHBOC’s) (**Figure S4**). Taken with the particle size results (**Figure 2A, F**), this suggests that particles formed with 90-minute extracted silk at 0.25% (w/v) will be predominantly dense polymeric particles without a distinct core. Thus, moving forward, we will only evaluate hemoglobin incorporation in this formulation (90 min-0.25%-X Hb) given the homogeneity along the radial direction in silk structure.

### 2.2. Introducing hemoglobin to silk fibroin nanoparticles

Knowing that, using aqueous solvent evaporation, silk particles can be formed in our target size range and that there is a substantial presence of β-sheet structures, we hypothesized that bioactive components such as hemoglobin could be kinetically trapped in silk secondary structures. This can be achieved by introducing the component to the silk prior to the hydrophobic collapse and structural formation of the biomaterial. This strategy for passive incorporation and stabilization of bioactive molecules and proteins has been investigated and reported previously for a variety of silk fibroin-based materials that contain β-sheet structures.^42, 54-56^ However, it is critical to confirm 3 main properties of our particle system with the introduction of hemoglobin: 1) limited change in size or polydispersity of the particles so that the neat silk fibroin particles are able to act as controls for future work 2) The spatial orientation of hemoglobin allow for better understanding of intraparticle homogeneity 3) the amount of incorporated hemoglobin that remains active and able to bind and release oxygen. To assess changes in particle size, morphology, and dispersity, we incorporate a ferric (non-oxygen binding) form of hemoglobin termed methemoglobin (mHb). We perform DLS and SEM in order to understand how addition of the mHb protein impacts the particle properties.

#### 2.2.1. Passive encapsulation of hemoglobin increases particles size, but maintains similar polydispersity

With the addition of hemoglobin, at either 125 or 250 μg of hemoglobin/mL of silk-PVA solution, particles size increases by 100-150 nm, taking average particle size of the 90-minute extracted silk at 0.25% silk solution from ~300 nm to ~400 nm (**Figure 4A**). However, the polydispersity of these formulations improved with the inclusion of mHb as shown by the peak width (**Figure 4B**). Taken together, **Figure 4AB** show that even though average particle size is increasing, the dispersity is such that for all initial mHb concentrations, there is significant overlap in the size of particles present. For neat silk particles the peak range given by the 68% maximum is 50-550 nm with an average of 300 nm, and for the 250 µg/mL of HbA_0_ included this shifts to a 250-650 nm range with a 450 average. This indicates that while we lose some of the smallest diameter particles from the population, there is still substantial overlap in the particle size for the tested formulations. These findings are also supported by the SEM micrographs for the respective conditions which show persistence of similarly sized particles across initial mHb concentrations (**Figure 4 CDE**). These results indicate that with future filtration strategies, a consistent size distribution can be maintained across levels of hemoglobin inclusion. We also performed similar analyses for other silk fibroin particle formulations and present those results in **Figure S4**.

**Figure 4:**
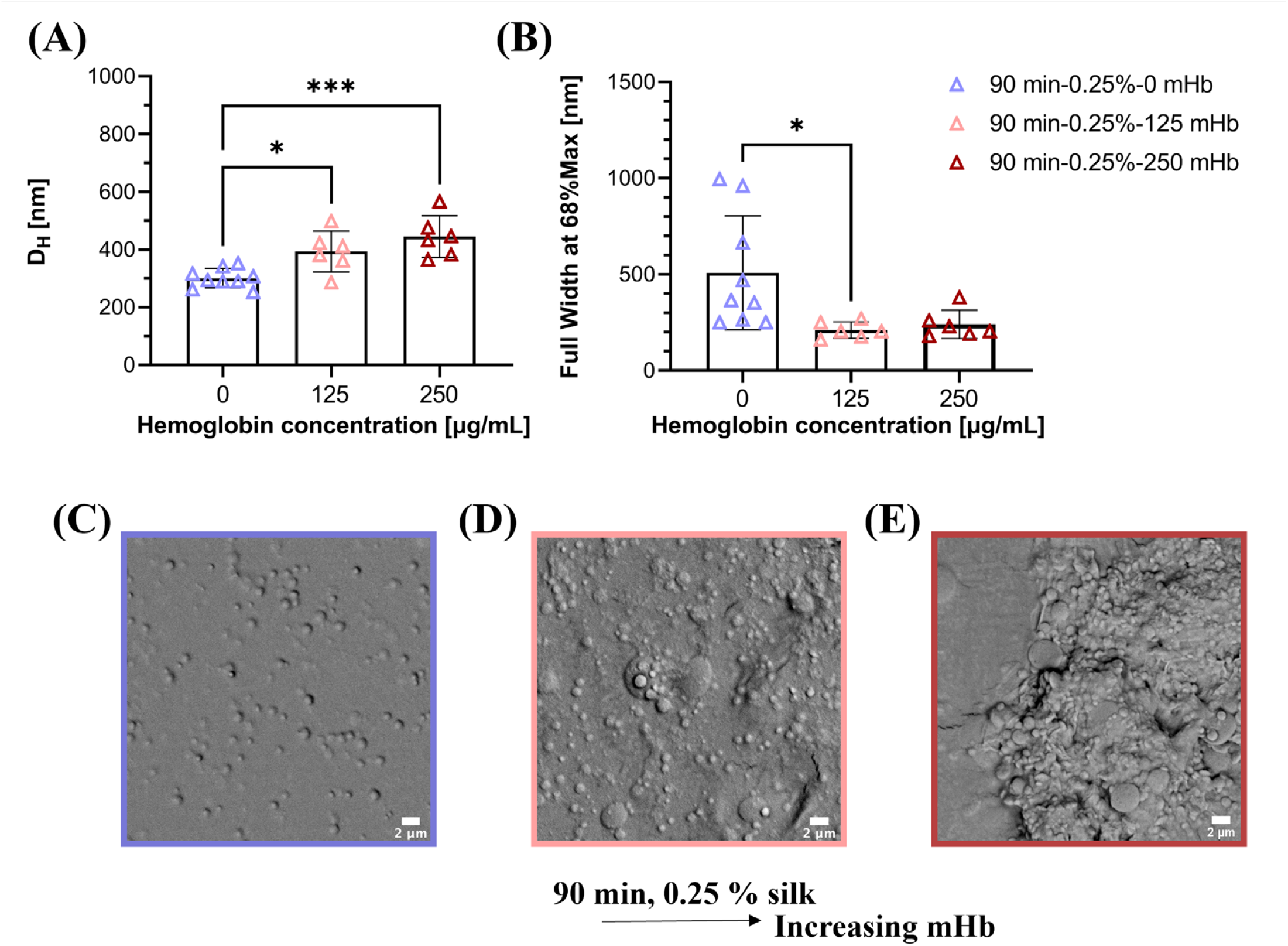
As hemoglobin concentration approaches silk concentration, average particle size increases slightly. (**A**) Comparison of hydrodynamic diameter (DH) for 90 min-0.25%-X formulations as measured by DLS. (**B**) Comparison of peak width at 68% of the maximum for 90 min-0.25%-X formulations, an assessment of dispersity. (**C**) Representative SEM micrograph for 90 min-0.25%-0 mHb (**D**) Representative SEM micrograph for 90 min-0.25%-125 mHb. (**E**) Representative SEM micrograph for 90 min-0.25%-250 mHb. Data shown are AVG ± SD (n≥6). A one-way ANOVA with Tukey post hoc testing was performed. Statistical significance is shown as *p<=0.05, **p<=0.01, ***p<=0.001, ****p<=0.0001.

#### 2.2.2. Passive encapsulation of functional human hemoglobin maintains O2 binding capacity

To confirm we encapsulated hemoglobin into the particle matrix, we quantified hemoglobin incorporation (**Figure 5 ABC**) and performed immunofluorescence microscopy to visualize hemoglobin incorporation (**Figure 5D**). Particles were made using a ferrous (oxygen binding) form of hemoglobin, referred to as HbA0. Following the same methods described in Section 2.2.1., we then used a FITC-conjugated antibody with a human hemoglobin A target. In **Figure 5D** we show colocalization of the hemoglobin signal (green) with the silk fibroin particles signal (blue^57^). We observe limited unspecific binding of the antibody (**Figure 5D, i**) and observe that with increasing initial HbA0 concentrations, Hb can agglomerate in the wall or on the edges of particles (**Figure 5D, iii**). When the silk is in excess relative to the hemoglobin, we observe disperse hemoglobin inclusion across the particle surface (**Figure 5D, ii**). Knowing that we can encapsulate mHb in the silk particle matrix, we then confirmed that our organic solvent free method maintains the gas binding capabilities of hemoglobin. To assess this, we encapsulated the ferrous form of hemoglobin (HbA_0_) in the same particle formations and utilized the cyanmethemoglobin method and spectrophotometry to analyze both the overall hemoglobin encapsulation efficiency as well as the levels of methemoglobin (ferric, non-active hemoglobin) present in the samples. We found that independent of the initial HbA_0_ concentration efficiency remained constant at approximately 60% (**Figure 5A**). This suggested that we were on the maximum side of the level of encapsulation we can achieve.

**Figure 5:**
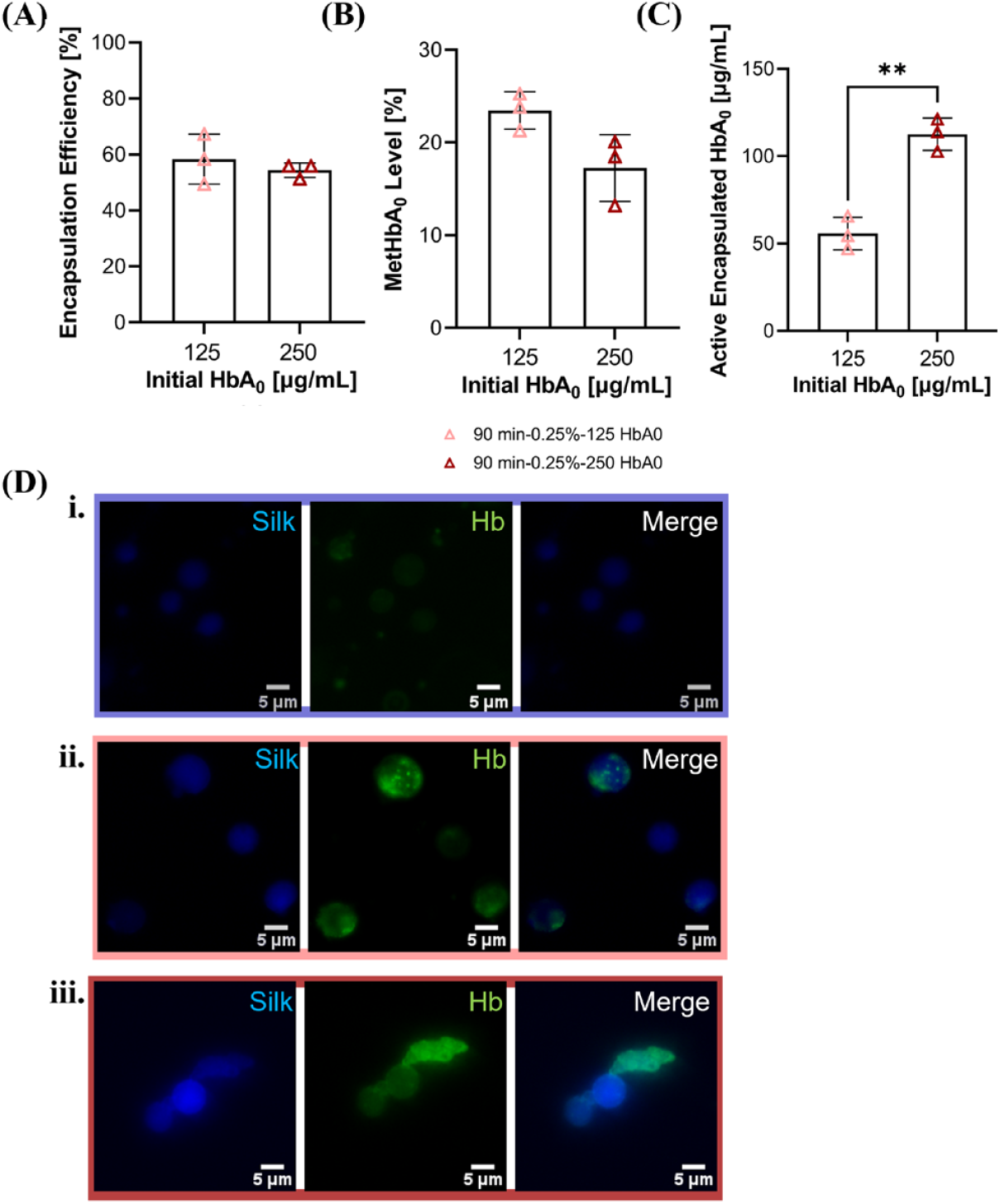
Functional hemoglobin is incorporated into silk particles. (**A**) Hemoglobin encapsulation efficiency was determined for 90 min-.25% (w/v) silk formulations with initial hemoglobin concentrations of 125 and 250 µg/mL using the cyanmethemoglobin method. (**B**) The level of methemoglobin (a oxidized, non-oxygen binding form of hemoglobin) was determined for 90 min-.25% (w/v) silk formulations with initial hemoglobin concentrations of 125 and 250 µg/mL using the cyanmethemoglobin method. (**C**) Combining the results shown in Figure 5 A and B, we calculated the concentration of active (oxygen binding) hemoglobin present for 90 min-.25% (w/v) silk formulations with initial hemoglobin concentrations of 125 and 250 µg/mL. (**D**) Fluorescent stain for hemoglobin in 90 min-1% (w/v) silk formulations with initial concentrations of (**i**) 0, (**ii**) 125 and, (**iii**) 250 µg/mL. silk is shown in blue and hemoglobin is shown in green. Data shown are AVG ± SD (n=3). A student t-test is performed. Statistical significance is shown as *p<=0.05, **p<=0.01.

To prove this, we performed the same experiment with increased initial HbA_0_ concentrations and found that encapsulation efficiency continues to decline for increasing initial HbA_0_ concentrations (**Figure S7A**). We then used the cyanmethemoglobin method to assess the level of methemoglobin in the samples (**Figures S6B and S7B**). From here, the amount of total active hemoglobin was calculated from the initial HbA_0_ concentration multiplied by the encapsulation efficiency, subtracting out the contribution of methemoglobin. We found that a maximum of 23% of encapsulated was in the ferric methemoglobin form (**Figure S7B**). Other research in the HBOC field has shown that methemoglobin levels around 10% allowed for substantial oxygenation of tissues.^58^ Native red blood cells in a healthy individual have levels of methemoglobin of only ~1% largely due to reductase enzymes allowing the methemoglobin to be reversed to its ferrous form. Some liposome based HBOCs have been able to nearly match the level of methemoglobin initially after formulation when freshly isolated hemoglobin from expired blood units are used.^21, 59^ We hypothesize that improved hemoglobin sourcing and timelines can further improve the level of methemoglobin we observe in our system. However, for applications not replacing more than 30% of the blood volume methemoglobin levels of ~20% in HBOCs can be helpful in supporting native red blood cells.^58^ We found that initial HbA^0^ concentrations of 250 µg/mL led to similar levels of ~100 µg/mL active hemoglobin in both 90-minute 0.25% and 90-minute 1% formulations (**Figure 5B and Figure S6D**). This is approximately 1/1000 the level of hemoglobin in a healthy individual, suggesting that these formulations could be used for applications where supplementation of red blood cells is necessary. These clinical situations include blood loss related to trauma in a non-clinical setting (*e*.*g*., battlefield, car accident), blood loss in immunocompromised patients where transfusion of human donor blood is sub-optimal (*e*.*g*., post childbirth, during a pandemic), or situations where a patient is unable to receive a transfusion due to religious convictions.

### 2.3. Silk particles do not illicit pro-inflammatory response in macrophages

Finally, to analyze the potential utility sfHBOCs, we investigated how macrophages, important phagocytes of the innate immune system, respond to sfHBOCs in culture. Understanding the ability of our particles to alter expression of key markers in mouse macrophage-like cells will give insight into future experimental design as we move toward clinical translation of silk fibroin particle technologies. Macrophages are an important cell type to understand the behavior of with this technology because resident macrophages of the liver and spleen are responsible for the clearance of cell-free Hb, and for the clearance of particles or any foreign matter from circulation. In homeostatic conditions, cell-free hemoglobin is cleared by liver macrophages via a CD163 mediated uptake of hemoglobin-haptoglobin complexes.^60-62^ In an ideal HBOC system, resident macrophages of the liver and spleen will be able to carry out this homeostatic function without being overwhelmed/stimulated by the carrier itself. Here we are describing a silk fibroin-based carrier, so it is important to understand the impact silk fibroin particles have on monocyte/macrophage-like cells. Implanted neat silk fibroin materials have shown evidence of innate immune phagocytes infiltration at the site of implantation for up to 2 weeks via analysis of CD68 expression, suggesting only mild and acute inflammation, an important step in the tissue healing cascade.^42, 63^ Previous studies with silk fibroin particles engineered for IP delivery also show mild inflammatory response by macrophages via the monitoring of TNF-α both pre- and post-translation.^48, 64, 65^ Here we observe limited activation of murine blood derived RAW 264.7 macrophages via the assessment of iNOS protein expression in cultures that are stimulated twice over 72 hours (**Figure 6**). This was observed for both neat silk particles as well as hemoglobin containing particles. Where when the percent of iNOS positive cells was normalized to the ground M0 control, all particle formulations resulted in a statistically similar decrease in iNOS expression and the only significant increase in iNOS positive cells was in the M1 positive control group. Additionally, if the percent of iNOS positive cells is normalized instead to the M1 control we observe all particle conditions, like the anti-inflammatory (M2) control and M0 control, maintain a negative fold change from the positive control group (**Figure S8**). Taken together this assay demonstrates the potential of silk particles to be cleared effectively and safely via native processes without substantially impacting long term macrophage phenotype or activity.

**Figure 6:**
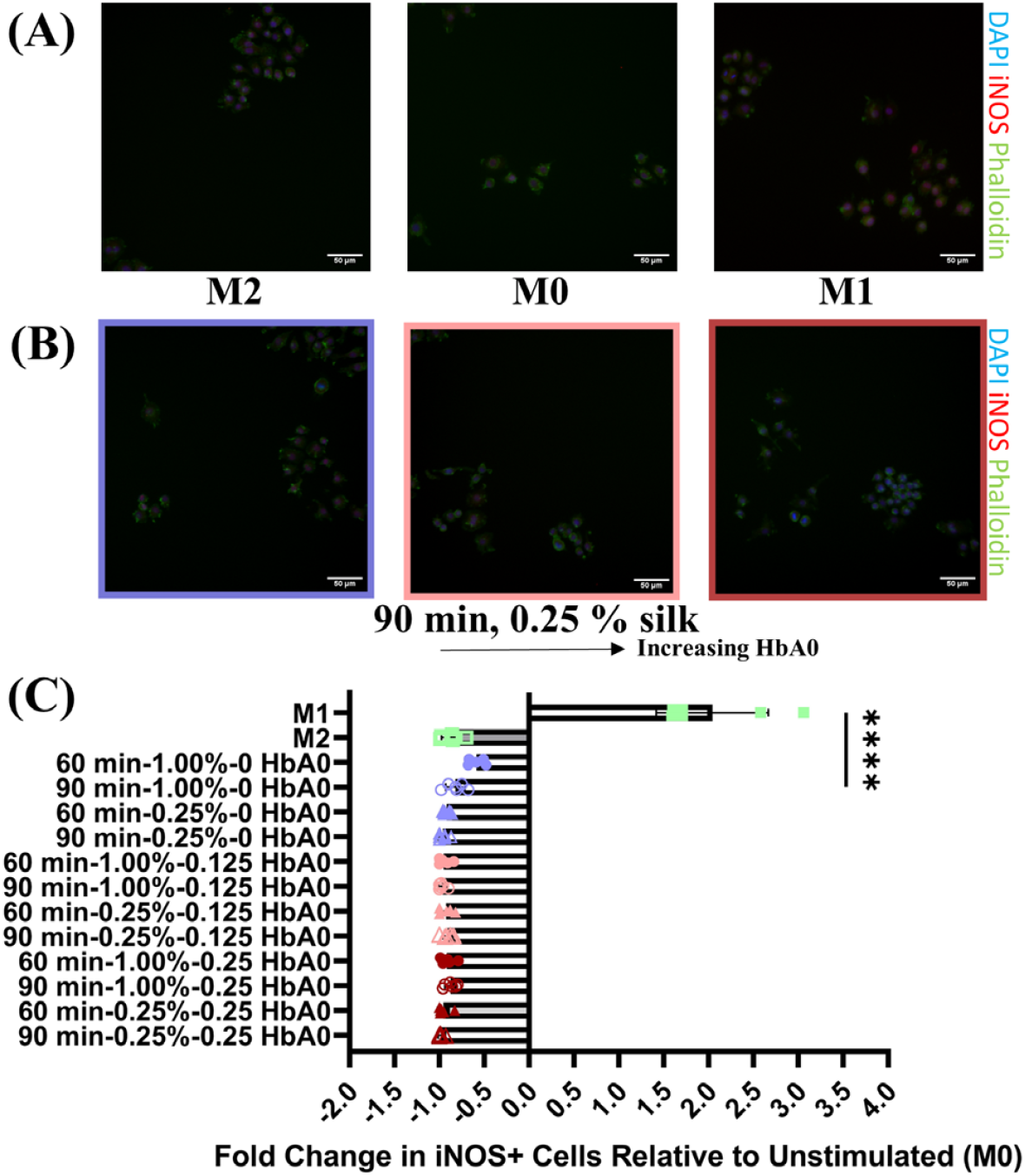
iNOS expression in RAW 264.7 is not significantly increased with particle incubation. (**A**) Flourescent images of negative (M2) ground (M0) and positive (M1) controls for iNOS (red) expression. Cells were also stained for nuclei (DAPI, blue), as well as spreading (Phalloidin, green) (**B**) Flourescent images for 90 min-0.25%-X HbA_0_ formulations. iNOS (red), DAPI (blue), Phalloidin (green) (**C**) Quantified fold change in iNOS expressing cells normalized to ground (M0) control. Data shown as AVG±SD (n=6). A one-way ANOVA with Tukey post hoc testing was performed. Statistical significance is shown as *p<=0.05, **p<=0.01, ***p<=0.001, ****p<=0.0001.

## 3. Conclusion

The results of this work have leveraged the properties of silk fibroin, an all-natural and non-thrombogenic biopolymer, to engineer a nanoparticle system as a carrier for hemoglobin, generating a silk fibroin-hemoglobin-based oxygen carrier (sfHBOC) as an alternative delivery vehicle to the liposome-dominated field of HBOCs. The results of this study demonstrate an aqueous processing method can yield control over particle size, achieving particles <500 nm in diameter with stable encapsulation of active hemoglobin (ability to bind O_2_ based on oxidation state) at levels needed for hemoglobin supplementation in clinical applications. These formulations do not elicit unwanted immune responses from macrophage-like cells *in vitro*.

Furthermore, the crystalline content in silk fibroin particles is critical for understanding the potential to include and stabilize hemoglobin within these particles to create sfHBOCs. Proteins and biomolecules are stabilized within the crystalline domains of the silk protein, trapping the bioactive molecule in a stable confirmation even when the particles experience various hydration levels or temperatures. This feature of silk fibroin biomaterials is what makes this all-natural carrier system advantageous as a HBOC.

In conclusion, the silk fibroin-hemoglobin-based oxygen carrier (sfHBOC) has the potential to address the toxicity and adverse reactions associated with commercially available HBOCs. These findings hold significant promise for the development of an injectable HBOC that can address the limitations of current HBOCs and provide effective oxygen delivery in transfusion medicine and the treatment of diseases. Further research is needed to evaluate the safety and efficacy of sfHBOCs *in vivo*.

## 4. Experimental Section

### 4.1. Silk solution preparation

Silk fibroin is isolated from *Bombyx mori* cocoons as previously described.^44^ Briefly, 5 grams of dime-size sections of silk cocoons are boiled to extract fibroin proteins from sericin proteins for 15, 30, 60, or 90 min in 2 L of 0.02 M aqueous sodium carbonate (Na_2_CO_3_) solution (Catalog No. 451614, Sigma Aldrich, St. Louis, MO). Once degumming is complete, the fibroin mat is air-dried for a minimum of 48 hours. The solid protein mat is then solubilized at 60 °C for 4 hours in 9.3 M aqueous lithium bromide (LiBr) solution (Catalog No. 213225, Sigma Aldrich, St. Louis, MO). This solution is then dialyzed against ultrapure water using 3.5 kDa MW cutoff dialysis membrane tubing (Spectrum™ Spectra/Por™ 3 RC Dialysis Membrane Tubing, 3,500 Dalton MWCO, Catalog No. 08-670-5C, ThermoFisher Scientific, Waltham, MA) over 48 hours to remove ions. The solubilized, aqueous silk solution is then centrifuged (4000 x g, 20 min, 4°C) twice to remove any insoluble impurities. The concentration (weight/volume) is determined by drying a known volume of the silk solution at 60°C and weighing the remaining solids. This protocol resulted in a 5–7% weight per volume (w/v) silk fibroin solution. Silk solutions were stored at 4°C for a maximum of 3 weeks prior to use in making particles.

### 4.2. Gel electrophoresis

Silk molecular weight (MW) distributions after fiber degumming were visualized with sodium dodecyl sulfate polyacrylamide gel electrophoresis (SDS-PAGE). Samples of varied extraction times (15, 30, 60, and 90 minutes) and 4 batches of 60-minute extracted silk were analyzed. For each sample 2 μg of solubilized silk protein was loaded into a NuPAGE™ 7% Tris-Acetate gel (EA03585BOX, Invitrogen, Waltham, MA) under reducing conditions. 5 μL of a high molecular weight ladder (HiMark™ Pre-stained Protein Standard LC5699, Invitrogen) was loaded in lane 1 of each gel. Gels were loaded into an Invitrogen mini gel tank (A25977) and run with Bolt™ 1X MES SDS Running buffer (Life Technologies, Carlsbad, CA). Using a power ease 300W (Life Technologies) power supply each gel was run at 100 V for 5 minutes and then the voltage was increased to 200 V for 23 minutes. The gels were stained with a Colloidal Blue staining kit (LC6025, Invitrogen) and then imaged on an Odyssey Fc (LI COR, Lincoln, NE) at 700 nm with a 30 second exposure time. The MW distributions of the solubilized silk samples were quantified with densitometric measurements along the length of each lane (Image Studio version 5.2, LI COR).

### 4.3. Silk and silk-hemoglobin particle preparation

Silk fibroin control particles are formed via a phase separation from polyvinyl alcohol (PVA) (P8136, Sigma Aldrich) modified from previous techniques.^44, 45^ Briefly, stock solutions of 5% (w/v) of silk and PVA are prepared. The silk and PVA are combined in a 1:4 weight ratio in a total volume of 5 mL. The resulting solution is then probe sonicated at 25% for 30 seconds. Post-sonication, the solution is cast into a 100 × 15 mm petri dish. Once dried overnight, the plastic-like film is shaken for 30 min in 20 mL dI water to dissolve. The resulting solution is centrifuged (4000 x g, 20 min, 4°C). The pellet containing silk particles is then resuspended in 5 mL of ultrapure water and sonicated at 15% for 15 seconds. In an effort to modulate the size of the particles, the extraction time of the silk used, and the silk concentration were varied as summarized in **Table 1**. A weight ratio of 1:4 silk:PVA was held constant across all conditions. Hemoglobin is passively encapsulated by combining the desired concentration of hemoglobin with silk solution prior to combining with PVA. For assessment of particle size and morphology bovine hemoglobin (H2625, Sigma Aldrich), which is primarily in the ferric form was used (referred to as methemoglobin (mHb) throughout manuscript). For encapsulation and cell experiments a human hemoglobin in ferrous form was used (H0267, Sigma Aldrich), referred to as HbA0 throughout manuscript.

### 4.4. Fourier Transform Infrared Spectroscopy (FTIR)

Quantification of the β-sheet content and other secondary protein structures in silk particles was performed using Fourier transform infrared spectroscopy (FTIR) analysis. Prior to FTIR analysis, particles were lyophilized at -80°C and 0.160 mbar for 2 days (FreeZone 12 Liter −84°C Console Freeze Dryer, Labconco, Kansas City, MO). Spectra were collected with a Nicolet iS50 FTIR Spectrometer (ThermoFisher Scientific, Waltham, MA) (UF Nanoscale Research Facility), equipped with a micro attenuated total reflections (microATR) germanium crystal and MCT/A detector. Each measurement consisted of 128 scans with a resolution of 4cm^-1^ over wavenumbers ranging 4,000-650 cm^-1^. Background spectra were collected using the same conditions and subtracted from each sample spectra. The amide I region (1,590–1,710 cm^− 1^) can be split into regions based on the protein secondary structure: 1,605–1,615 cm^−1^ as side chain/aggregated strands, 1,616–1,637 cm^−1^ and 1,697–1,703 cm^−1^ as β-sheet structure, 1,638–1,655 cm^−1^ as random coils, 1,656–1,662 cm^−1^ as α-helical bands, and 1,663–1,696 cm^−1^ as turns. For analysis the amide I region was deconvoluted to obtain relative amounts of respective secondary structures as detailed by Hu et. al.^1^

### 4.5. Nanoscale Computed Tomography (Nano-CT)

Assessment of internal particle morphology and detection of dense crystalline regions of silk was done through use of Nano-CT at the UF Nanoscale Research Facility. Particles we prepared using 60-minute extracted silk at 1% (w/v). The resulting particle solution was lyophilized (FreeZone 12 Liter -84°C Console Freeze Dryer, Labconco, Kansas City, MO) for 48 hours. The lyophilized particles were then imaged (v|tome|x m 240; General Electric) and analyzed using VGStudio Max.

### 4.6. Dynamic Light Scattering (DLS)

The particle size and poly dispersity of silk nanoparticle formulations in ultrapure water was determined at 25°C by dynamic light scattering (DLS) using a Zetasizer Nano-ZS Malvern Instrument. The refractive indices of 1.33 for water and 1.60 for silk protein are used as has been done previously for silk materials and all measurements are conducted in triplicate (error not propagated).^43^ In analysis of the dispersity of particle solutions, we determined the peak width at 68% lower than the peak maximum. This was done to represent a standard deviation since in a normal distribution 68% of the population falls within one standard deviation.^66^

### 4.7. Scanning Electron Microscopy (SEM)

10 µL of particle suspension in water was directly added to a ZEISS/LEO SEM Pin Stub Mount with an adhesive sticker, Ø12.7mm x 9mm pin height (Catalog No. 16202, Ted Pella, Inc., Redding, CA). The samples were then dried in a chemical hood for 48 hours and sputter-coated with 10 nm of Au prior to imaging on a Phenom Pure benchtop SEM.

### 4.8. Estimation of encapsulation efficiency and methemoglobin level by UV-Vis Spectroscopy

Spectrophotometric absorbance measurements were obtained using a NanoDrop One/One^c^ (ThermoFisher Scientific, Waltham MA). The total Hb and metHb (i.e., oxidized Hb in the Fe3+ valence state) concentrations were determined using the cyanmethemoglobin method.^67, 68^ To determine the total concentration of hemoglobin, 20% potassium ferricyanide K3(Fe(CN)6) (ThermoFisher Scientific, AC211095000) was added to samples and allowed to react for 3 min, converting all hemoglobin to methemoglobin. Then, 10% KCN (ThermoFisher Scientific, AAL1327322) is added to form cyanmethemoglobin with unique absorbance behavior. The absorbance was then measured at 540 nm. The final hemoglobin concentration is calculated as a dilution coefficient multiplied with the ratio of the absorbance and the extinction coefficient of methemoglobin at 540 nm [cm mM]^-1^ (**Figure S5A**). The methemoglobin levels of encapsulated hemoglobin were found using cyanmethemoglobin method and UV-vis.^67, 68^ The absorbance of the sample was measured at 630 nm. Then a 1:1 KCN and PBS solution was added to samples. This converts methemoglobin to cyanomethemoglobin, which does not absorb at 630 nm. After a minimum of 3 minutes, the absorbance was read again at 630 nm. The concentration of methemoglobin can be calculated as a dilution coefficient multiplied by the ratio of the change in absorbance over the extinction coefficient for methemoglobin at 630 nm [cm mM]^-1^ (**Figure 5B**).

### 4.9. Fluorescent imaging of hemoglobin

Hemoglobin encapsulation in silk particles was visualized using fluorescent microscopy. Samples prepared with 90 min-1% silk and initial HbA_0_ (H0267, Sigma Aldrich) concentrations of 0, 125, and 250 µg/mL. Following particle formation, 100 µL of particle solution was placed in microcentrifuge tubes and combined with blocking media (5% donkey serum in 1% bovine serum albumin (BSA) in PBS) and incubated at room temperature for a minimum of 4 hours. After blocking, wells were washed 3 times with PBS with 0.1% tween (PBST) and then incubated overnight at 4 °C with a human hemoglobin-FITC conjugated antibody (Life Technologies, A80134F). Samples were washed three times with PBST prior to imaging on a Keyence BZ-X800 microscope.

### 4.10. RAW 264.7 culture

The murine BALB/c derived RAW 264.7 cell line was obtained from ATCC (Manassas, Va). The cells maintained in Dulbecco’s Modified Eagle’s Medium (DMEM) (Gibco) which was supplemented with 10% fetal bovine serum (FBS) (Gibco) and 1% Penicillin-Streptomycin (15-140-122, Gibco). This base media formulation serves as the M0 control media. Cells were routinely subcultured every 3-4 days and maintained at 37 °C in 5% CO2. Macrophage activations was assessed by seeding at 1.05 × 10^4^ cells/cm^2^ and allowing them to recover overnight. M1 stimulation media was M0 control media with 10 ng/ml IFNγ (Peprotech) and 100 ng/ml Lipopolysaccharide (LPS) (SigmaAldrich) added. M2 stimulation media was M0 media with 40 ng/ml IL-4 (Peprotech) and 20 ng/ml IL-13 (Peprotech) added. Stimulation with M1 media, M2 media, and silk particles occurred 24 hours after seeding. 250 μl of stimulation media was added to each well with unstimulated wells receiving M0 media. The wells stimulated with silk nanoparticles received 250 μl of M0 media supplemented with 50 μg of silk particles. 48 hours post-stimulation the media was changed using the same conditions as the first stimulation. This stimulation strategy resulted in 15 groups 72 hours post-stimulation: M0 (unstimulated), M1-Like, M2-Like, and the 12 silk nanoparticle formulations.

### 4.11. Immunostaining

An immunofluorescence assay was conducted to analyze the expression of iNOS (M1 marker) and cell spreading following the stimulation of RAW 264.7 cells. The 72 hours post-stimulation cells were fixed in 10% Phosphate buffered formalin for 15 minutes at room temperature and then washed 3 times with phosphate buffered saline (PBS). The cells were then permeabilized with 0.05% Triton-X-100 for 20 minutes at room temperature and washed 3 times with PBS. Following permeabilization each well was blocked for 5 hours with 5% donkey serum in 1% bovine serum albumin (BSA) in PBS at room temperature. After blocking, wells were washed 3 times with PBS with 0.1% tween (PBST) and then incubated overnight at 4 °C with the anti-iNOS primary antibody (Rabbit polyclonal Antibody, PA5-1624, Invitrogen) at 0.002 ng/ml in 1% BSA in PBS. Secondary control wells were incubated with only 1% BSA in PBS. Following primary incubation wells were washed 3 times with PBST prior to incubation with the anti-rabbit secondary antibody (A31572, Invitrogen) at 2.5 μg/ml and phalloidin at 16.7 μg/ml for 2 hours at room temperature in the dark. DAPI (ThermoFisher Scientific) was added to each well at 5 μg/ml for 10 min in the dark. Wells were washed with PBST 3 times before adding 200 μl PBST to each well and fluorescent microscopy was conducted with the Keyence BZ-X800 microscope. iNOS expression in images was quantified using Cell Profiler™.

### 4.12. Statistical methods

Experimental data are mainly expressed as mean ± standard deviation (SD) with a minimum n=3. In the case of the gel electrophoresis data, data are expressed as median ± half of the inner quartile range (**Figure 1**). GraphPad Prism 8.4.1 (La Jolla, CA) was utilized to analyze these data. Analysis is completed with appropriate-sized analysis of variance (ANOVA). If significance was found, Tukey post hoc testing was used for pairwise comparisons. Statistical significance is reported as *p < 0.05, **p < 0.01, ***p < 0.001, and ****p < 0.0001, with key features shown in the figures and full information provided in the supplement. Specifics on the statistical test, p values, and definition of n are all present in individual figure captions.

## Supporting information

Pacheco_SupplementalMaterials

Pacheco_SupplementalVideo1

## 5. Acknowledgements and Funding

The Stoppel lab would like to acknowledge the research scientists at the University of Florida Nanoscale Research Facility, especially Gary Scheiffele, PhD, for their assistance with using core equipment, such as the Nano-CT and FTIR. Additionally, support from Stoppel Lab University of Florida undergraduates in silk solution preparation, including Travis Truong, Hannah Bagnis, and Marina Fernandez-Campa. We also appreciate assistance from Dr. Yeongseon Jang and her PhD student Jackson Powers for the use of their Malvern DLS instrument.

All authors would like to acknowledge support from the Department of Defense Congressionally Directed Medical Research Fund (W81XWH2110199). ASC acknowledges support from the Chemical Engineering REU Site at the University of Florida (NSF EEC-1852111). NAD was supported by a grant from the National Institutes of Health (NIH T35HL007489). MOP acknowledges support from the National Science Foundation Graduate Research Fellowship (DGE-1842473). Any opinions, findings, and conclusions or recommendations expressed in this manuscript are those of the authors and do not necessarily reflect the views of the National Science Foundation, National Institutes of Health, or Department of Defense.

## 6. Conflict of Interest

Whitney L. Stoppel, Bruce D. Spiess, and Marisa O. Pacheco have a pending patent on this research.

## References

1. Hu X, Kaplan D, Cebe P. Determining beta-sheet crystallinity in fibrous proteins by thermal analysis and infrared spectroscopy. Macromolecules. 2006;39(18):6161–70. doi: 10.1021/ma0610109. PubMed PMID: WOS:000240069300032.

2. Dumont LJ, Cancelas JA, Maes LA, Rugg N, Whitley P, Herschel L, Siegal AH, Szczepiorkowski ZM, Hess JR, Zia M. Overnight, room temperature hold of whole blood followed by 42-day storage of red blood cells in additive solution-7. Transfusion. 2015;55(3):485–90. Epub 20140919. doi: 10.1111/trf.12868. PubMed PMID: 25234026.

3. Czubak K, Antosik A, Cichon N, Zbikowska HM. Vitamin C and Trolox decrease oxidative stress and hemolysis in cold-stored human red blood cells. Redox Rep. 2017;22(6):445–50. Epub 20170211. doi: 10.1080/13510002.2017.1289314. PubMed PMID: 28277068; PMCID: PMC6837575.

4. Zou S. Potential impact of pandemic influenza on blood safety and availability. Transfus Med Rev. 2006;20(3):181–9. Epub 2006/06/22. doi: 10.1016/j.tmrv.2006.03.001. PubMed PMID: 16787826; PMCID: PMC7134961.

5. Gilcher RO, McCombs S. Seasonal blood shortages can be eliminated. Curr Opin Hematol. 2005;12(6):503–8. Epub 2005/10/12. doi: 10.1097/01.moh.0000180436.98990.ce. PubMed PMID: 16217170.

6. Haracic M, Simmonds S, Krausbauer U. Maintaining blood supply during war and siege conclusions from experience in Sarajevo, Bosnia-Herzegovina. Transfusion Medicine and Hemotherapy. 2003;30(1):37–9. doi: Doi 10.1159/000069343. PubMed PMID: WOS:000182597400005.

7. Miskeen E, Omer Yahia AI, Eljack TB, Karar HK. The Impact of COVID-19 Pandemic on Blood Transfusion Services: A Perspective from Health Professionals and Donors. J Multidiscip Healthc. 2021;14:3063–71. Epub 20211102. doi: 10.2147/JMDH.S337039. PubMed PMID: 34754194; PMCID: PMC8572088.

8. Fahimnia B, Jabbarzadeh A, Ghavamifar A, Bell M. Supply chain design for efficient and effective blood supply in disasters. International Journal of Production Economics. 2017;183:700–9. doi: 10.1016/j.ijpe.2015.11.007. PubMed PMID: WOS:000391899700007.

9. Sen Gupta A. Hemoglobin-based Oxygen Carriers: Current State-of-the-art and Novel Molecules. Shock. 2019;52(1S Suppl 1):70–83. Epub 2019/09/13. doi: 10.1097/SHK.0000000000001009. PubMed PMID: 31513123; PMCID: PMC6874912.

10. Lane TA. Perfluorochemical-based artificial oxygen carrying red cell substitutes. Transfus Sci. 1995;16(1):19–31. Epub 1995/02/07. doi: 10.1016/0955-3886(94)00067-t. PubMed PMID: 10155703.

11. Ketcham EM, Cairns CB. Hemoglobin-based oxygen carriers: development and clinical potential. Ann Emerg Med. 1999;33(3):326–37. doi: 10.1016/s0196-0644(99)70370-7. PubMed PMID: 10036348.

12. Manning JE, Katz LM, Brownstein MR, Pearce LB, Gawryl MS, Baker CC. Bovine hemoglobin-based oxygen carrier (HBOC-201) for resuscitation of uncontrolled, exsanguinating liver injury in swine. Carolina Resuscitation Research Group. Shock. 2000;13(2):152–9. Epub 2000/02/12. doi: 10.1097/00024382-200013020-00010. PubMed PMID: 10670846.

13. Sakai H, Masada Y, Takeoka S, Tsuchida E. Characteristics of bovine hemoglobin as a potential source of hemoglobin-vesicles for an artificial oxygen carrier. J Biochem. 2002;131(4):611–7. Epub 2002/04/03. doi: 10.1093/oxfordjournals.jbchem.a003141. PubMed PMID: 11927000.

14. Cao M, Zhao Y, He H, Yue R, Pan L, Hu H, Ren Y, Qin Q, Yi X, Yin T, Ma L, Zhang D, Huang X. New Applications of HBOC-201: A 25-Year Review of the Literature. Front Med (Lausanne). 2021;8:794561. Epub 20211208. doi: 10.3389/fmed.2021.794561. PubMed PMID: 34957164; PMCID: PMC8692657.

15. Meng F, Kassa T, Jana S, Wood F, Zhang X, Jia Y, D’Agnillo F, Alayash AI. Comprehensive Biochemical and Biophysical Characterization of Hemoglobin-Based Oxygen Carrier Therapeutics: All HBOCs Are Not Created Equally. Bioconjug Chem. 2018;29(5):1560–75. Epub 20180402. doi: 10.1021/acs.bioconjchem.8b00093. PubMed PMID: 29570272.

16. Sarin H. Physiologic upper limits of pore size of different blood capillary types and another perspective on the dual pore theory of microvascular permeability. J Angiogenes Res. 2010;2:14. Epub 20100811. doi: 10.1186/2040-2384-2-14. PubMed PMID: 20701757; PMCID: PMC2928191.

17. Alayash AI. Mechanisms of Toxicity and Modulation of Hemoglobin-based Oxygen Carriers. Shock. 2019;52(1S Suppl 1):41–9. Epub 2017/11/08. doi: 10.1097/SHK.0000000000001044. PubMed PMID: 29112106; PMCID: PMC5934345.

18. Vallelian F, Buehler PW, Schaer DJ. Hemolysis, free hemoglobin toxicity, and scavenger protein therapeutics. Blood. 2022;140(17):1837–44. doi: 10.1182/blood.2022015596. PubMed PMID: 35660854.

19. Jahr JS, Akha AS, Holtby RJ. Crosslinked, polymerized, and PEG-conjugated hemoglobin-based oxygen carriers: clinical safety and efficacy of recent and current products. Curr Drug Discov Technol. 2012;9(3):158–65. Epub 2011/07/13. doi: 10.2174/157016312802650742. PubMed PMID: 21745179.

20. Kuang L, Zhu Y, Wu Y, Tian K, Peng X, Xue M, Xiang X, Lau B, Tzang FC, Liu L, Li T. A Novel Cross-Linked Hemoglobin-Based Oxygen Carrier, YQ23, Extended the Golden Hour for Uncontrolled Hemorrhagic Shock in Rats and Miniature Pigs. Front Pharmacol. 2021;12:652716. Epub 20210512. doi: 10.3389/fphar.2021.652716. PubMed PMID: 34054533; PMCID: PMC8149754.

21. Cuddington CT, Wolfe SR, Belcher DA, Allyn M, Greenfield A, Gu X, Hickey R, Lu S, Salvi T, Palmer AF. Pilot scale production and characterization of next generation high molecular weight and tense quaternary state polymerized human hemoglobin. Biotechnol Bioeng. 2022;119(12):3447–61. Epub 20221003. doi: 10.1002/bit.28233. PubMed PMID: 36120842; PMCID: PMC9828582.

22. Muller CR, Williams AT, Walser C, Eaker AM, Sandoval JL, Cuddington CT, Wolfe SR, Palmer AF, Cabrales P. Safety and efficacy of human polymerized hemoglobin on guinea pig resuscitation from hemorrhagic shock. Sci Rep. 2022;12(1):20480. Epub 20221128. doi: 10.1038/s41598-022-23926-y. PubMed PMID: 36443351; PMCID: PMC9703428.

23. Farcas AD, Toma VA, Roman I, Sevastre B, Scurtu F, Silaghi-Dumitrescu R. Glutaraldehyde-Polymerized Hemoglobin: In Search of Improved Performance as Oxygen Carrier in Hemorrhage Models. Bioinorg Chem Appl. 2020;2020:1096573. Epub 20200901. doi: 10.1155/2020/1096573. PubMed PMID: 32952540; PMCID: PMC7482000.

24. Kawaguchi AT, Fukumoto D, Haida M, Ogata Y, Yamano M, Tsukada H. Liposome-encapsulated hemoglobin reduces the size of cerebral infarction in the rat: evaluation with photochemically induced thrombosis of the middle cerebral artery. Stroke. 2007;38(5):1626–32. Epub 20070329. doi: 10.1161/STROKEAHA.106.467290. PubMed PMID: 17395856.

25. Azuma H, Fujihara M, Sakai H. Biocompatibility of HbV: Liposome-Encapsulated Hemoglobin Molecules-Liposome Effects on Immune Function. J Funct Biomater. 2017;8(3):24. Epub 20170628. doi: 10.3390/jfb8030024. PubMed PMID: 28657582; PMCID: PMC5618275.

26. Rameez S, Palmer AF. Simple method for preparing poly(ethylene glycol)-surface-conjugated liposome-encapsulated hemoglobins: physicochemical properties, long-term storage stability, and their reactions with O2, CO, and NO. Langmuir. 2011;27(14):8829–40. Epub 20110616. doi: 10.1021/la201246m. PubMed PMID: 21678920; PMCID: PMC3148852.

27. Kure T, Sakai H. Preparation of Artificial Red Blood Cells (Hemoglobin Vesicles) Using the Rotation-Revolution Mixer for High Encapsulation Efficiency. ACS Biomater Sci Eng. 2021;7(6):2835–44. Epub 20210524. doi: 10.1021/acsbiomaterials.1c00424. PubMed PMID: 34029046.

28. Awasthi V, Yadav VR, Goins B, Phillips WT. Modulation of oxidative stability of haemoglobin inside liposome-encapsulated haemoglobin. J Microencapsul. 2013;30(5):471–8. Epub 20121211. doi: 10.3109/02652048.2012.752535. PubMed PMID: 23231644; PMCID: PMC3696053.

29. Akbarzadeh A, Rezaei-Sadabady R, Davaran S, Joo SW, Zarghami N, Hanifehpour Y, Samiei M, Kouhi M, Nejati-Koshki K. Liposome: classification, preparation, and applications. Nanoscale Res Lett. 2013;8(1):102. Epub 20130222. doi: 10.1186/1556-276X-8-102. PubMed PMID: 23432972; PMCID: PMC3599573.

30. Lopez RR, P GFdR, Sanchez LM, Tsering T, Alazzam A, Bergeron KF, Mounier C, Burnier JV, Stiharu I, Nerguizian V. The effect of different organic solvents in liposome properties produced in a periodic disturbance mixer: Transcutol(R), a potential organic solvent replacement. Colloids and surfaces B, Biointerfaces. 2021;198:111447. Epub 20201104. doi: 10.1016/j.colsurfb.2020.111447. PubMed PMID: 33223347.

31. Bandyopadhyay A, Chowdhury SK, Dey S, Moses JC, Mandal BB. Silk: A Promising Biomaterial Opening New Vistas Towards Affordable Healthcare Solutions. J Indian I Sci. 2019;99(3):445–87. doi: 10.1007/s41745-019-00114-y. PubMed PMID: WOS:000491592800012.

32. Kluge JA, Li AB, Kahn BT, Michaud DS, Omenetto FG, Kaplan DL. Silk-based blood stabilization for diagnostics. Proc Natl Acad Sci U S A. 2016;113(21):5892–7. Epub 20160509. doi: 10.1073/pnas.1602493113. PubMed PMID: 27162330; PMCID: PMC4889389.

33. Li AB, Kluge JA, Guziewicz NA, Omenetto FG, Kaplan DL. Silk-based stabilization of biomacromolecules. J Control Release. 2015;219:416–30. Epub 20150925. doi: 10.1016/j.jconrel.2015.09.037. PubMed PMID: 26403801; PMCID: PMC4656123.

34. Lee KY, Kong SJ, Park WH, Ha WS, Kwon IC. Effect of surface properties on the antithrombogenicity of silk fibroin/S-carboxymethyl kerateine blend films. J Biomater Sci Polym Ed. 1998;9(9):905–14. doi: 10.1163/156856298x00235. PubMed PMID: 9747984.

35. Maitz MF, Sperling C, Wongpinyochit T, Herklotz M, Werner C, Seib FP. Biocompatibility assessment of silk nanoparticles: hemocompatibility and internalization by human blood cells. Nanomedicine. 2017;13(8):2633–42. Epub 20170727. doi: 10.1016/j.nano.2017.07.012. PubMed PMID: 28757180.

36. Hu X, Kaplan D, Cebe P. Dynamic Protein−Water Relationships during β-Sheet Formation. Macromolecules. 2008;41(11):3939–48. doi: 10.1021/ma071551d.

37. Matthew SAL, Totten JD, Phuagkhaopong S, Egan G, Witte K, Perrie Y, Seib FP. Silk Nanoparticle Manufacture in Semi-Batch Format. ACS Biomater Sci Eng. 2020;6(12):6748–59. Epub 20201103. doi: 10.1021/acsbiomaterials.0c01028. PubMed PMID: 33320640.

38. Totten JD, Wongpinyochit T, Carrola J, Duarte IF, Seib FP. PEGylation-Dependent Metabolic Rewiring of Macrophages with Silk Fibroin Nanoparticles. ACS Appl Mater Interfaces. 2019;11(16):14515–25. Epub 20190412. doi: 10.1021/acsami.8b18716. PubMed PMID: 30977355.

39. Florczak A, Deptuch T, Lewandowska A, Penderecka K, Kramer E, Marszalek A, Mackiewicz A, Dams-Kozlowska H. Functionalized silk spheres selectively and effectively deliver a cytotoxic drug to targeted cancer cells in vivo. J Nanobiotechnology. 2020;18(1):177. Epub 20201201. doi: 10.1186/s12951-020-00734-y. PubMed PMID: 33261651; PMCID: PMC7709326.

40. Montalban MG, Coburn JM, Lozano-Perez AA, Cenis JL, Villora G, Kaplan DL. Production of Curcumin-Loaded Silk Fibroin Nanoparticles for Cancer Therapy. Nanomaterials (Basel). 2018;8(2):126. Epub 20180224. doi: 10.3390/nano8020126. PubMed PMID: 29495296; PMCID: PMC5853757.

41. Pacheco MO, Eccles LE, Davies NA, Armada J, Cakley AS, Kadambi IP, Stoppel WL. Progress in silk and silk fiber-inspired polymeric nanomaterials for drug delivery. Frontiers in Chemical Engineering. 2022;4.

42. Jameson JF, Pacheco MO, Bender EC, Kotta NM, Black LD, Kaplan DL, Grasman JM, Stoppel WL. Impact of bioactive molecule inclusion in lyophilized silk scaffolds varies between in vivo and in vitro assessments. bioRxiv. 2022:2022.05.24.493207. doi: 10.1101/2022.05.24.493207.

43. Wongpinyochit T, Johnston BF, Seib FP. Manufacture and Drug Delivery Applications of Silk Nanoparticles. J Vis Exp. 2016(116):e54669. Epub 20161008. doi: 10.3791/54669. PubMed PMID: 27768078; PMCID: PMC5092179.

44. Rockwood DN, Preda RC, Yucel T, Wang X, Lovett ML, Kaplan DL. Materials fabrication from Bombyx mori silk fibroin. Nat Protoc. 2011;6(10):1612–31. Epub 20110922. doi: 10.1038/nprot.2011.379. PubMed PMID: 21959241; PMCID: PMC3808976.

45. Wang X, Yucel T, Lu Q, Hu X, Kaplan DL. Silk nanospheres and microspheres from silk/pva blend films for drug delivery. Biomaterials. 2010;31(6):1025–35. Epub 20091127. doi: 10.1016/j.biomaterials.2009.11.002. PubMed PMID: 19945157; PMCID: PMC2832579.

46. Wray LS, Hu X, Gallego J, Georgakoudi I, Omenetto FG, Schmidt D, Kaplan DL. Effect of processing on silk-based biomaterials: reproducibility and biocompatibility. J Biomed Mater Res B Appl Biomater. 2011;99(1):89–101. Epub 20110621. doi: 10.1002/jbm.b.31875. PubMed PMID: 21695778; PMCID: PMC3418605.

47. Partlow BP, Tabatabai AP, Leisk GG, Cebe P, Blair DL, Kaplan DL. Silk Fibroin Degradation Related to Rheological and Mechanical Properties. Macromol Biosci. 2016;16(5):666–75. Epub 20160112. doi: 10.1002/mabi.201500370. PubMed PMID: 26756449.

48. Montoya NV, Peterson R, Ornell KJ, Albrecht DR, Coburn JM. Silk Particle Production Based on silk/PVA Phase Separation Using a Microfabricated Co-flow Device. Molecules. 2020;25(4). Epub 20200217. doi: 10.3390/molecules25040890. PubMed PMID: 32079339; PMCID: PMC7070425.

49. Rahmani H, Fattahi A, Sadrjavadi K, Khaledian S, Shokoohinia Y. Preparation and Characterization of Silk Fibroin Nanoparticles as a Potential Drug Delivery System for 5-Fluorouracil. Adv Pharm Bull. 2019;9(4):601–8. Epub 20191024. doi: 10.15171/apb.2019.069. PubMed PMID: 31857964; PMCID: PMC6912188.

50. Matthew SAL, Rezwan R, Perrie Y, Seib FP. Volumetric Scalability of Microfluidic and Semi-Batch Silk Nanoprecipitation Methods. Molecules. 2022;27(7):2368. Epub 20220406. doi: 10.3390/molecules27072368. PubMed PMID: 35408763; PMCID: PMC9000471.

51. Wongpinyochit T, Johnston BF, Seib FP. Degradation Behavior of Silk Nanoparticles-Enzyme Responsiveness. ACS Biomater Sci Eng. 2018;4(3):942–51. Epub 20180220. doi: 10.1021/acsbiomaterials.7b01021. PubMed PMID: 33418776.

52. Crivelli B, Perteghella S, Bari E, Sorrenti M, Tripodo G, Chlapanidas T, Torre ML. Silk nanoparticles: from inert supports to bioactive natural carriers for drug delivery. Soft Matter. 2018;14(4):546–57. Epub 2018/01/13. doi: 10.1039/c7sm01631j. PubMed PMID: 29327746.

53. Solomun JI, Totten JD, Wongpinyochit T, Florence AJ, Seib FP. Manual Versus Microfluidic-Assisted Nanoparticle Manufacture: Impact of Silk Fibroin Stock on Nanoparticle Characteristics. ACS Biomater Sci Eng. 2020;6(5):2796–804. Epub 20200420. doi: 10.1021/acsbiomaterials.0c00202. PubMed PMID: 32582839; PMCID: PMC7304816.

54. Stoppel WL, Hu D, Domian IJ, Kaplan DL, Black LD, 3rd. Anisotropic silk biomaterials containing cardiac extracellular matrix for cardiac tissue engineering. Biomedical materials (Bristol, England). 2015;10(3):034105. Epub 20150331. doi: 10.1088/1748-6041/10/3/034105. PubMed PMID: 25826196; PMCID: PMC4417360.

55. Reeves AR, Spiller KL, Freytes DO, Vunjak-Novakovic G, Kaplan DL. Controlled release of cytokines using silk-biomaterials for macrophage polarization. Biomaterials. 2015;73:272–83. Epub 20150921. doi: 10.1016/j.biomaterials.2015.09.027. PubMed PMID: 26421484; PMCID: PMC4605898.

56. Stinson JA, Palmer CR, Miller DP, Li AB, Lightner K, Jost H, Weldon WC, Oberste MS, Kluge JA, Kosuda KM. Thin silk fibroin films as a dried format for temperature stabilization of inactivated polio vaccine. Vaccine. 2020;38(7):1652–60. Epub 20200117. doi: 10.1016/j.vaccine.2019.12.062. PubMed PMID: 31959422; PMCID: PMC7176408.

57. Amirikia M, Shariatzadeh SMA, Jorsaraei SGA, Mehranjani MS. Auto-fluorescence of a silk fibroin-based scaffold and its interference with fluorophores in labeled cells. Eur Biophys J. 2018;47(5):573–81. Epub 20180212. doi: 10.1007/s00249-018-1279-1. PubMed PMID: 29435602.

58. Linberg R, Conover CD, Shum KL, Shorr RG. Hemoglobin based oxygen carriers: how much methemoglobin is too much? Artif Cells Blood Substit Immobil Biotechnol. 1998;26(2):133–48. doi: 10.3109/10731199809119772. PubMed PMID: 9564432.

59. Banerjee U, Wolfe S, O’Boyle Q, Cuddington C, Palmer AF. Scalable production and complete biophysical characterization of poly(ethylene glycol) surface conjugated liposome encapsulated hemoglobin (PEG-LEH). PLoS One. 2022;17(7):e0269939. Epub 20220708. doi: 10.1371/journal.pone.0269939. PubMed PMID: 35802716; PMCID: PMC9269976.

60. Buehler PW, Abraham B, Vallelian F, Linnemayr C, Pereira CP, Cipollo JF, Jia Y, Mikolajczyk M, Boretti FS, Schoedon G, Alayash AI, Schaer DJ. Haptoglobin preserves the CD163 hemoglobin scavenger pathway by shielding hemoglobin from peroxidative modification. Blood. 2009;113(11):2578–86. Epub 20090108. doi: 10.1182/blood-2008-08-174466. PubMed PMID: 19131549.

61. Munoz CJ, Pires IS, Baek JH, Buehler PW, Palmer AF, Cabrales P. Apohemoglobin-haptoglobin complex attenuates the pathobiology of circulating acellular hemoglobin and heme. American journal of physiology Heart and circulatory physiology. 2020;318(5):H1296–H307. Epub 20200417. doi: 10.1152/ajpheart.00136.2020. PubMed PMID: 32302494; PMCID: PMC7346542.

62. Nielsen MJ, Andersen CB, Moestrup SK. CD163 binding to haptoglobin-hemoglobin complexes involves a dual-point electrostatic receptor-ligand pairing. J Biol Chem. 2013;288(26):18834–41. Epub 20130513. doi: 10.1074/jbc.M113.471060. PubMed PMID: 23671278; PMCID: PMC3696659.

63. Thurber AE, Omenetto FG, Kaplan DL. In vivo bioresponses to silk proteins. Biomaterials. 2015;71:145–57. Epub 20150820. doi: 10.1016/j.biomaterials.2015.08.039. PubMed PMID: 26322725; PMCID: PMC4573254.

64. Wongpinyochit T, Uhlmann P, Urquhart AJ, Seib FP. PEGylated Silk Nanoparticles for Anticancer Drug Delivery. Biomacromolecules. 2015;16(11):3712–22. Epub 20151013. doi: 10.1021/acs.biomac.5b01003. PubMed PMID: 26418537.

65. Totten JD, Wongpinyochit T, Carrola J, Duarte IF, Seib FP. PEGylation-Dependent Metabolic Rewiring of Macrophages with Silk Fibroin Nanoparticles. ACS Applied Materials & Interfaces. 2019;11(16):14515–25. doi: 10.1021/acsami.8b18716.

66. Cumming G, Williams J, Fidler F. Replication and Researchers’ Understanding of Confidence Intervals and Standard Error Bars. Understanding Statistics. 2004;3(4):299–311. doi: 10.1207/s15328031us0304_5.

67. Whitehead RD, Jr., Mei Z, Mapango C, Jefferds MED. Methods and analyzers for hemoglobin measurement in clinical laboratories and field settings. Ann N Y Acad Sci. 2019;1450(1):147–71. Epub 20190604. doi: 10.1111/nyas.14124. PubMed PMID: 31162693; PMCID: PMC6709845.

68. Rameez S, Alosta H, Palmer AF. Biocompatible and biodegradable polymersome encapsulated hemoglobin: a potential oxygen carrier. Bioconjug Chem. 2008;19(5):1025–32. Epub 20080429. doi: 10.1021/bc700465v. PubMed PMID: 18442283.

