## Supplementary material for "Silk Fibroin Particles as Carriers in the Development of All-Natural Hemoglobin-Based Oxygen Carriers (HBOCs)": Pacheco_SupplementalMaterials

**Author contact information:**

Marisa O Pacheco:
Henry M Lutz:
Jostin Armada:
Nickolas Davies:
Isabelle K Gerzenshtein:
Alaura S Cakley:

1. **Use of SDS Page Gel Electrophoresis to investigate average molecular weight of regenerated silk fibroin biopolymers in water**

When deciding the fabrication parameters most likely to modulate silk particle size, silk polymer length or molecular weight was identified as a main parameter of interest. To assess silk chain length a SDS-PAGE gel was run under reducing conditions. The gel was stained with a Colloidal Blue kit as described in the methods section of the manuscript and representative raw images of the stained gels for extraction time comparisons as well as batch-to-batch analysis are pictured here (**Figure S1**). Using a LiCOR Odyssey Imaging system, the gels shown in **Figure S1** were subjected to densiometric analysis, this yielded the results shown in **Figure 1** of the main manuscript.

2. **Representative analyses of particle size using dynamic light scattering**

To analyze particle size, Dynamic Light scattering was used. In the main body of the manuscript, we present the average intensity weighted hydrodynamic diameters for all neat silk particle samples. To gain a better understanding of the size distributions for these samples we first calculated the difference in the intensity reported average and the peak max. We expect that in relatively monodisperse populations, with particle sizes in the submicron range, for this difference to be small. We observe that 90-minute extracted silk-maintained agreement much better than 60-minute extracted silk (**Figure S2A**). This led us to focus distribution analysis to only 90-minute extracted conditions. The conclusion to focus distribution analysis determined by DLS on 90-minute extracted samples was also supported by the SEM micrographs shown in the main manuscript, in which 60-minute extracted samples show high levels of polydispersity and presence of larger particles. Below are representative intensity distributions and auto-correlation functions (ACF) for all silk only conditions. In any case where the ACF did not show an exponential decay shape

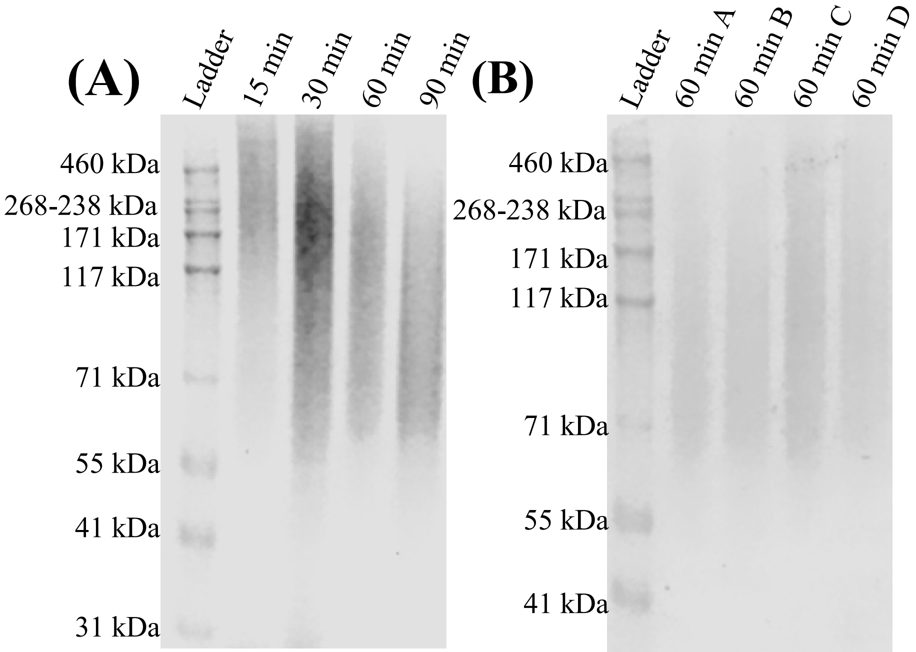


**Figure S1:** Representative raw images of Colloidal Blue stained SDS-PAGE gels for silk molecular weight analysis. (**A**) Comparing extraction times ranging from 15-90 minutes. (**B**) Comparing four separate 60 minute extractions for batch-to-batch analysis.

like those shown in Figure S2, the data point was not used, and measurements were repeated (**Figure S2 BCDE**).

### 3. FTIR as a method to interrogate secondary structures in silk nanoparticles

Fourier transform infrared (FTIR) spectroscopy was used to analyze the types of protein secondary structures present in the formed neat silk particles. In the main manuscript, we report the percent of β-crystalline structures present as they are central to silk fibroins ability to stabilize biomacromolecules. Here we report the levels of other protein secondary structures determined by deconvolution the Amide I region of the same captured spectra from **Figure 3A** (**Figure S3**). It is shown that higher silk concentration formulations (1% w/v silk) tend to show increased presence of random coil and α-helice content. This is complimentary to the trends observed in β-sheet content discussed in the main manuscript.

### 4. Nano-Computed Topography as a way to analyze internal structure of the silk particles

Nano-CT was used to assess the internal structure of the silk particles. In **Supplementary Video 1** we show a clipping plane moving through a 3D reconstruction of a silk microparticle. For this same particle we obtained line profiles of signal intensity or gray value, and this is reported in the main manuscript (**Figure 3C).** The line profiles indicate that there is less polymer density in the internal volume of the sphere, this video visually supports that conclusion. It also shows a polymer dense outer wall of the sphere. Analysis of wall thickness as it relates to particle size is also discussed in the main manuscript (**Figure 3DE**).

### 5. Impact of hemoglobin addition method on particle properties and hemoglobin structure

Upon the addition of mHb to neat silk formulations, we analyze the impact of Hb inclusion of the particle size and morphology. In the main manuscript we focus on showing the size and morphology results for the formulation that we hypothesize yielded the most uniform and radially symmetric particles (90 min, 0.25% silk). Herein, are the results for the other silk formulations when increasing amounts of hemoglobin are incorporated. Similar trends are observed to those presented in the main manuscript (**Figure S4**).

To determine the level of Hb encapsulation and the ability of that Hb to bind oxygen, we used the cyanmethemoglobin method with UV-vis spectroscopy. In the main manuscript we focus on the results for formulations using 90 min silk at 0.25%. Here we present details of how UV-Vis can be used to capture the encapsulation efficiency (**Figure S5A**) and level of methemoglobin (**Figure S5B**). Additionally, we show the results for 90-1% (w/v) silk formulations (**Figure S6**). Lastly, we show that at initial hemoglobin concentrations in excess of silk concentration, encapsulation is not efficient (**Figure S7**).

To better understand the potential safety and clearance of sfHBOCs we introduced them to RAW 264.7 cultures and assessed polarization. In the main manuscript, we present the results of staining for a proinflammatory (M1) marker iNOS normalized to M0 expression. Here, we show the same experiment normalized to the M1 positive control (**Figure S8**). This confirms that all conditions maintain a negative fold change difference from the positive control (M1) independent of how the normalization is done.

HU, X., KAPLAN, D. & CEBE, P. 2006. Determining Beta-Sheet Crystallinity in Fibrous Proteins by Thermal Analysis and Infrared Spectroscopy. *Macromolecules,* 39**,** 6161-6170.

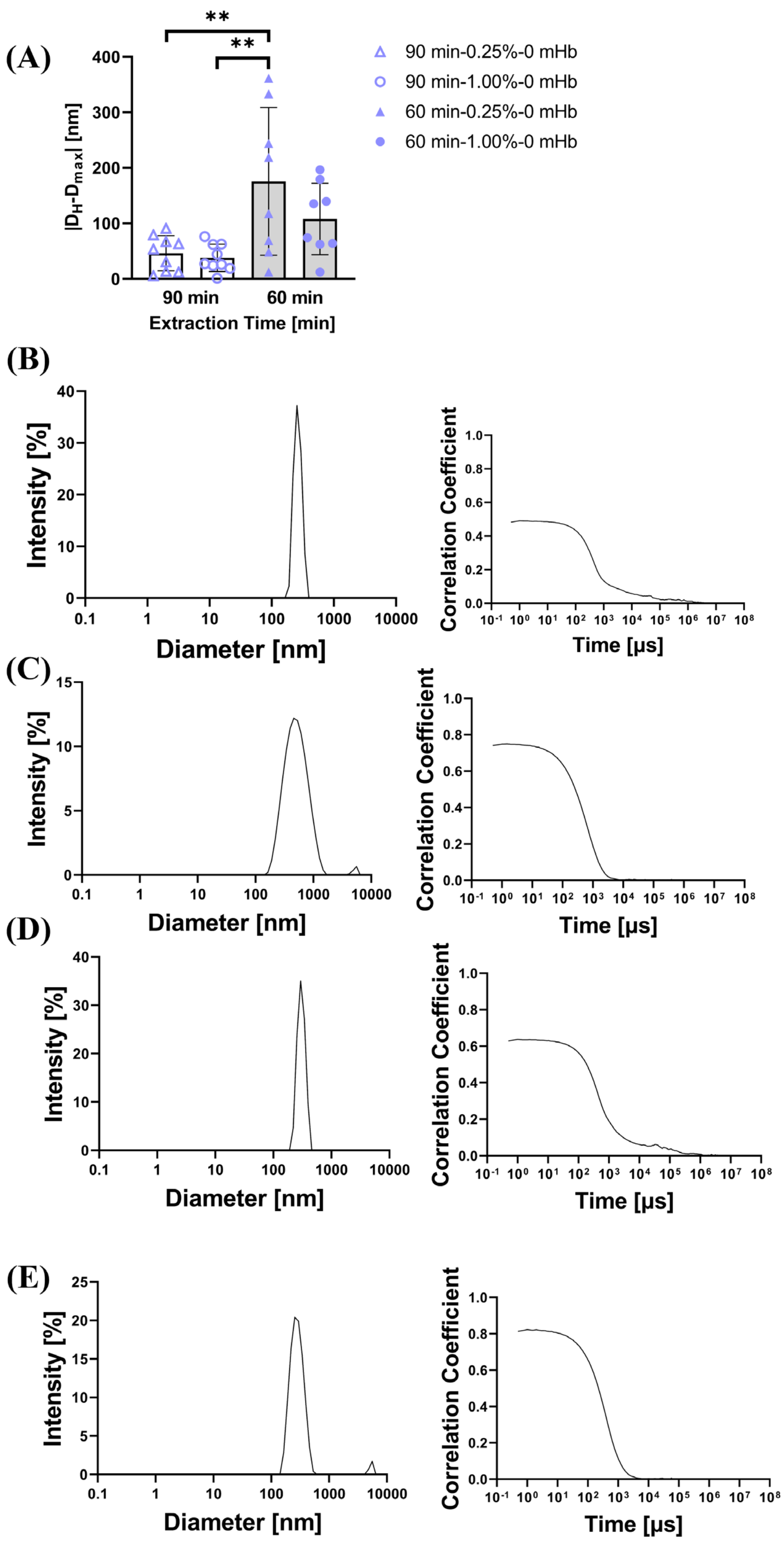


**Figure S2:** (**A**) The absolute value of the difference between the DLS determined intensity weighted average hydrodynamic diameter and the peak maximum diameter. Representative intensity size distributions (left) and autocorrelation functions (right) to accompany the DLS intensity size weighted mean particle hydrodynamic diameters presented in Figure 2 of the manuscript. (**B**) 60 min-1%-0Hb (**C**) 90min-1%-0Hb (**D**) 60 min-0.25%-0Hb (**E**) 90 min-0.25%-0Hb

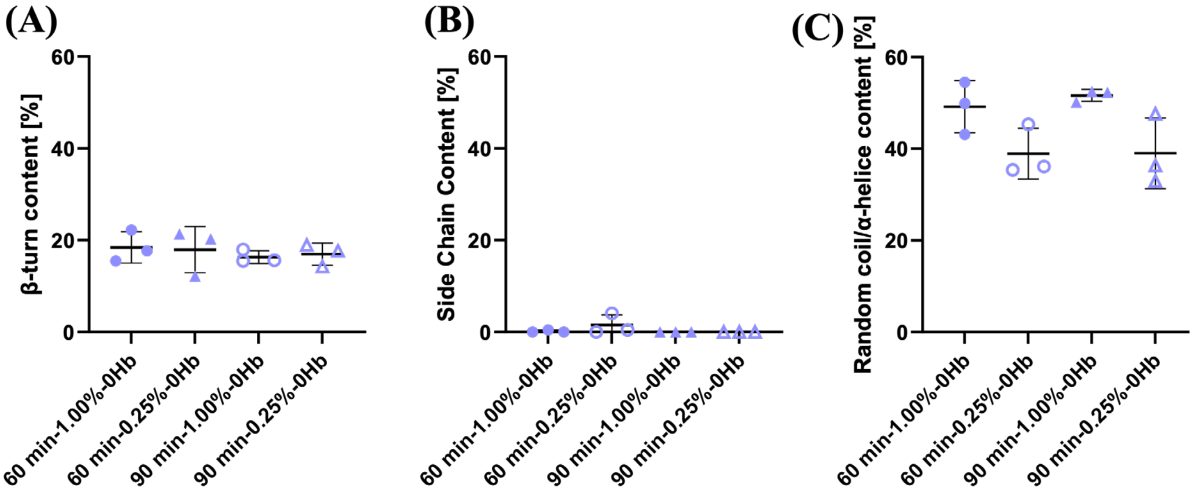


**Figure S3:** Estimated content of protein secondary structures based on the deconvolution of the Amide I region of captured FTIR spectra following methods previously described (Hu et al., 2006) (**A**) β-turns (**B**) Side chains (**C**) Random coils and α-helices. Data shown represent the average ± SD with n=3.

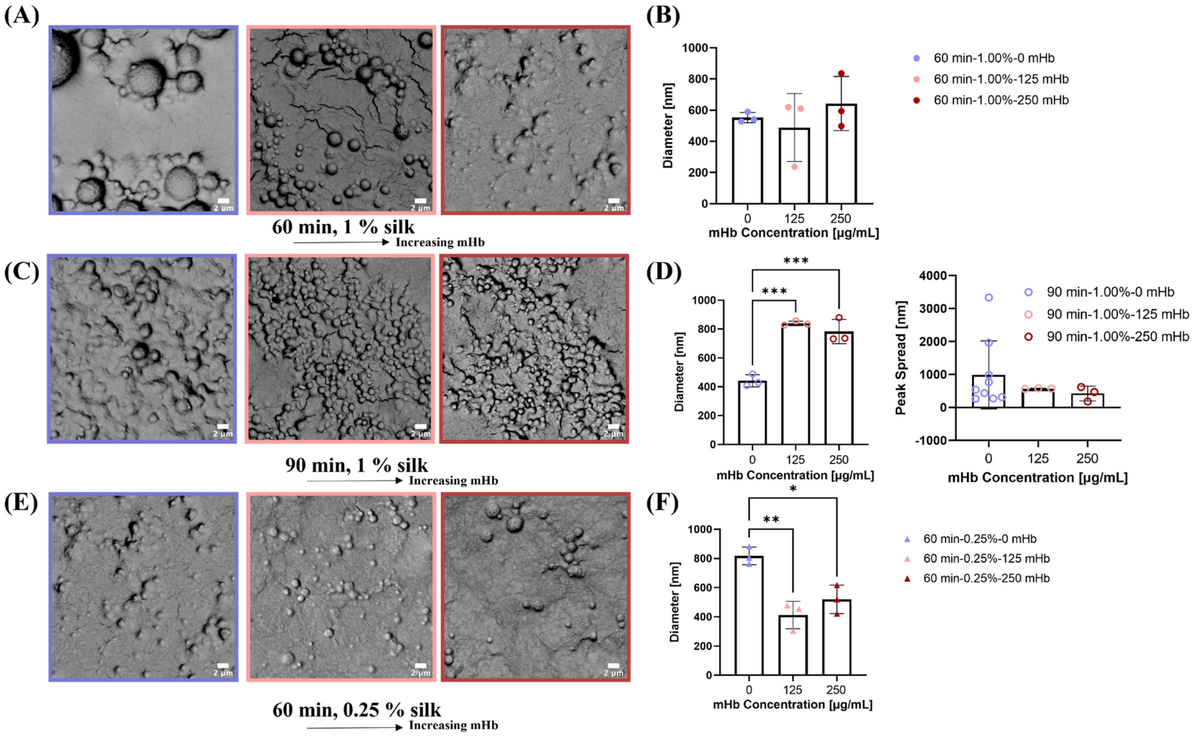


**Figure S4:** DLS and SEM Data for all conditions grouped as a function of neat silk formulation showing the impact of increasing hemoglobin inclusion. (**A**) Representative SEM micrographs of particles made with 60 min, 1% silk and varying levels of mHb. (**B**) Plots showing the intensity based average particle hydrodynamic for particles made with 60 min, 1% silk and varying levels of mHb. (**C**) Representative SEM micrographs of particles made with 90 min, 1% silk and varying levels of mHb. (**D**) Plots showing the intensity based average particle hydrodynamic diameter (left) and peak width (right) for particles made with 90 min, 1% silk and varying levels of mHb. (**E**) Representative SEM micrographs of particles made with 60 min, 0.25% silk and varying levels of mHb. (**F**) Plots showing the intensity based average particle hydrodynamic for particles made with 60 min, 0.25% silk and varying levels of mHb.

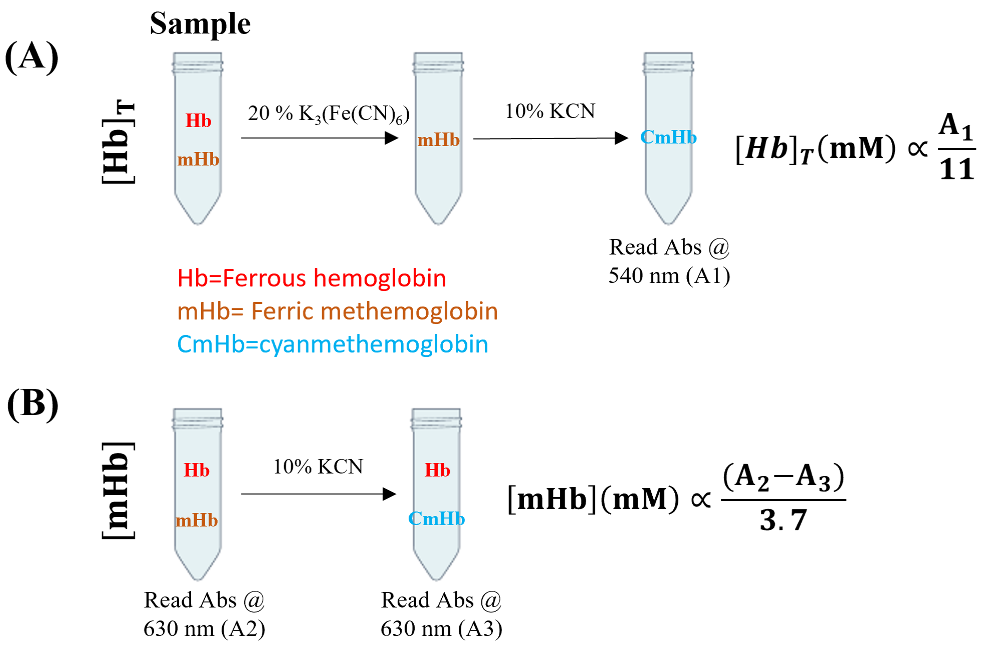


**Figure S5:** Cyanmethemoglobin method for determining hemoglobin concentrations. (**A**) To determine the total concentration of hemoglobin, 20% potassium ferricyanide $K_3(Fe(CN)_6)$ was added to samples and allowed to react for 3 min, converting all hemoglobin to methemoglobin. Then, 10% KCN is added to form cyanmethemoglobin with unique absorbance behavior. The absorbance was then measured at 540 nm (A1). The final hemoglobin concentration is calculated as a dilution coefficient multiplied with the ratio of the absorbance and the extinction coefficient of methemoglobin at 540 nm $[\text{cm mM}]^{-1}$. (**B**) The methemoglobin levels of encapsulated hemoglobin were found using cyanmethemoglobin method and UV-vis. The absorbance of the sample was measured at 630 nm (A2). Then a 1:1 potassium cyanide and PBS solution was added to samples. This converts methemoglobin to cyanomethemoglobin, which does not absorb at 630 nm. After a minimum of 3 minutes, the absorbance was read again at 630 nm (A3). The concentration of methemoglobin can be calculated as a dilution coefficient multiplied by the ratio of the change in absorbance over the extinction coefficient for methemoglobin at 630 nm $[\text{cm mM}]^{-1}$.

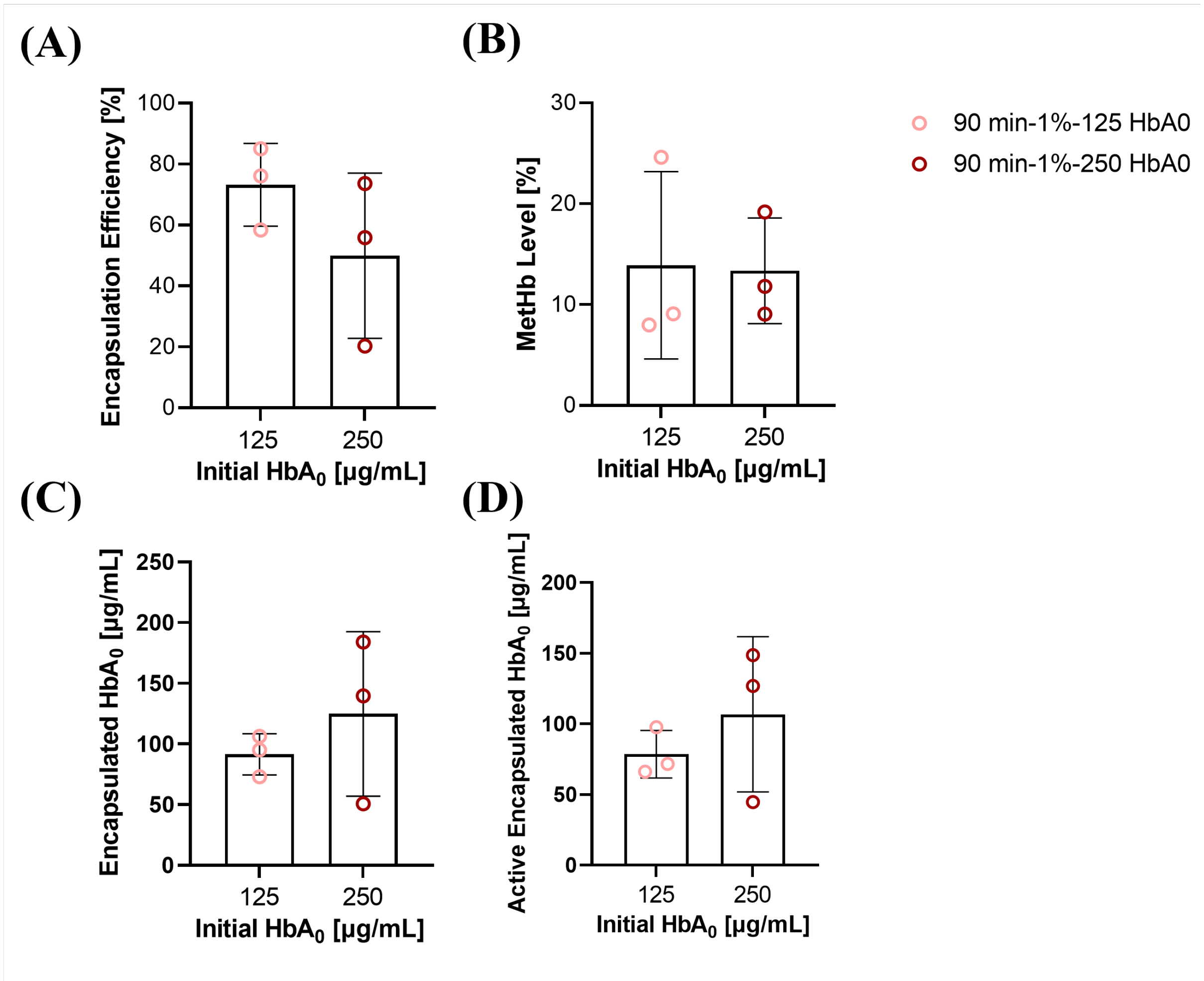


**Figure S6**: (**A**) Hemoglobin encapsulation efficiency was determined for 90 min-1% (w/v) silk formulations with initial hemoglobin concentrations of 125 and 250 µg/mL using the cyanmethemoglobin method. (**B**) The level of methemoglobin (a oxidized, non-oxygen binding form of hemoglobin) was determined for 90 min-1% (w/v) silk formulations with initial hemoglobin concentrations of 125 and 250 µg/mL using the cyanmethemoglobin method. (**C**) The total concentration of hemoglobin was calculated using the same methods that give encapsulation efficiency. (**D**) Combining the results shown in (A) and (B), we calculated the concentration of active (oxygen binding) hemoglobin present for 90 min-1% (w/v) silk formulations with initial hemoglobin concentrations of 125 and 250 µg/mL.

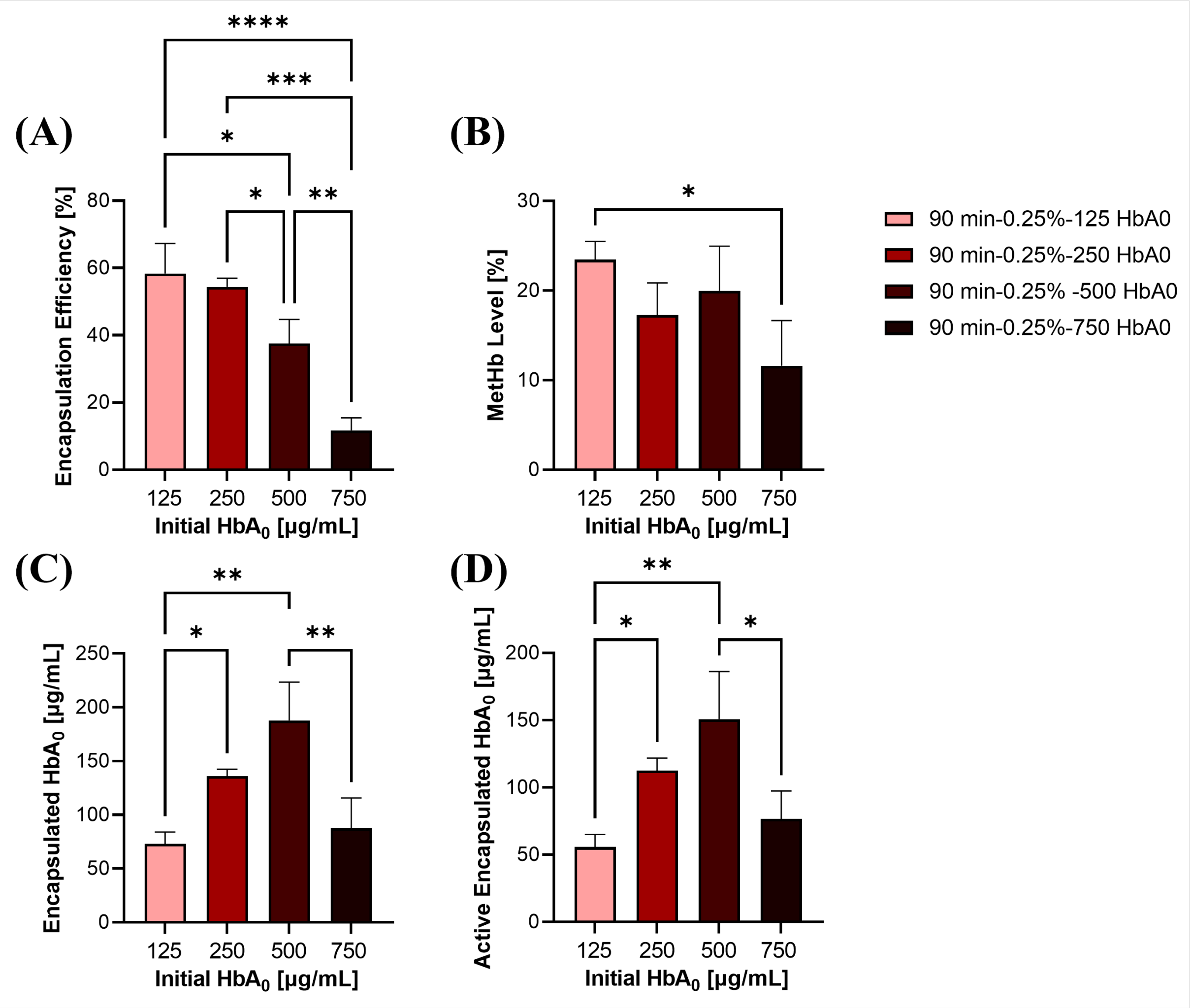


**Figure S7:** (**A**) Hemoglobin encapsulation efficiency was determined for 90 min-0.25% (w/v) silk formulations with initial hemoglobin concentrations of 125, 250, 500 and 750 µg/mL using the cyanmethemoglobin method. (**B**) The level of methemoglobin (a oxidized, non-oxygen binding form of hemoglobin) was determined for 90 min-0.25% (w/v) silk formulations with initial hemoglobin concentrations of 125, 250, 500 and 750 µg/mL using the cyanmethemoglobin method. (**C**) The total concentration of hemoglobin was calculated using the same methods that give encapsulation efficiency. (**D**) Combining the results shown in (A) and (B), we calculated the concentration of active (oxygen binding) hemoglobin present for 90 min-0.25% (w/v) silk formulations with initial hemoglobin concentrations of 125, 250, 500 and 750 µg/mL.

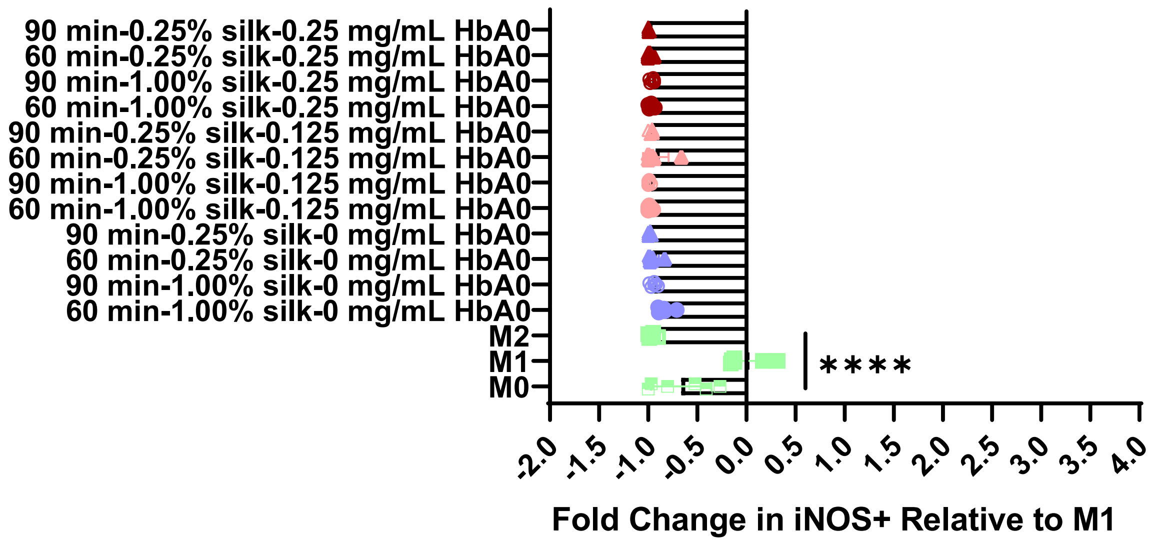


**Figure S8**: iNOS expression of RAW 264.7 macrophages following 3 days of exposure to silk particle formulations. Here we show the fold change in iNOS+ cells relative to a proinflammatory (M1) positive control (LPS, IFN-γ). Data are represented as mean±SD with n=6. A one way ANOVA was performed with Tukey post hoc testing. Statistical Significance is shown as ****p≤0.0001
